# Genome based analysis of Antibacterial Biosynthetic Clusters in *Lactiplantibacillus plantarum* C6 and Exploration of their Natural Small Molecules as anti-biofilm in Methicillin-Resistant *Staphylococcus aureus*

**DOI:** 10.1101/2025.09.04.674170

**Authors:** Daraksha Iram, Manish Singh Sansi, Ariel Fontana, Sudershan Kumar

**Affiliations:** Antimicrobial Peptides, Bio functional Probiotics & Peptidomics Laboratory, Dairy Microbiology Division, National Dairy Research Institute, Karnal, Haryana, India; Biofunctional Peptidomics & Metabolic Syndrome Laboratory, Animal Biochemistry Division, National Dairy Research Institute, Karnal, Haryana, India; ^e^Instituto de Hortofruticultura Subtropical y Mediterránea “La Mayora” (IHSM “La Mayora”), Universidad de Málaga-Consejo Superior de Investigaciones Científicas (UMA-CSIC), Campus Teatinos, 29010 Málaga, Spain; Grupo de Bioquímica Vegetal, Instituto de Biología Agrícola de Mendoza CONICET-UNCuyo, Almirante Brown 500, Chacras de Coria, M5507, Argentina; Cell Biology and Proteomics Lab, Animal Biotechnology Center, National Dairy Research Institute, Karnal-132001, India

**Keywords:** probiotics, metabolites, antimicrobials, lanthipeptides, adhesins, phylogeny, genome, probiotic, MRSA

## Abstract

This study presents the complete genome characterization of *Lactiplantibacillus plantarum* C6, a strain isolated from Indian dairy cheese, using Illumina NovaSeq sequencing. The assembled genome (3.22 Mb, 44.5% GC) comprised 3,076 coding sequences, 59 tRNAs, 10 rRNAs, and 2 CRISPR arrays. Phylogenomic and ANI analyses confirmed its identity within the *L. plantarum* clade (>99% similarity with NMGL2 and DMDL 9010). Functional annotation revealed genes enriched in carbohydrate metabolism (10.7%), stress response, and host-adaptation pathways, supporting its probiotic potential. Bacteriocin biosynthetic gene clusters were identified, including those encoding PlnE, PlnF, PlnJ, PlnK, and PlnN, indicating the strain’s ability to produce class II plantaricins. A RiPP cluster encoding a cyclic uberolysin-like peptide was also detected, with structural similarity to known lanthipeptides such as Streptococcin A, Nisin Q, and Lacticin 3147 (Tanimoto scores 0.93–1.0), suggesting antimicrobial relevance. CAZy analysis revealed 102 carbohydrate-active enzymes (GHs, GTs), highlighting metabolic flexibility. To evaluate the antibiofilm potential of *L. plantarum*-derived metabolites, 15 small molecules from cell-free supernatants (CFS) were selected through literature mining and subjected to molecular docking against the MRSA biofilm-associated enzyme poly-β-1,6-N-acetyl-D-glucosamine synthase (encoded by *icaA*). 2,4-Di-tert-butylphenol (−7.2 kcal/mol) and Indole-3-lactic acid (−7.1 kcal/mol) showed the strongest binding, followed by Cyclo (L-propyl-L-valine) (−6.8 kcal/mol) and DL-4-Hydroxyphenyllactic acid (−6.4 kcal/mol), indicating promising inhibition of MRSA biofilm synthesis. Organic acids like acetic and lactic acid showed weaker interactions but may contribute synergistically through acidification. Overall, *L. plantarum* C6 combines robust probiotic features, genomic safety, and antimicrobial potential, supported by bacteriocin gene clusters and effective antibiofilm metabolites, highlighting its application in functional foods and novel antimicrobial development.

## 1. Introduction

LAB play a multifaceted role in food matrices. They metabolize complex macromolecules, such as indigestible polysaccharides and off-flavour compounds, while generating beneficial metabolites: vitamins, biogenic amines, short-chain fatty acids, exopolysaccharides, and a variety of bacteriocins (Dawwam et al., 2022). Microbial contamination represents a major source of foodborne illnesses, posing persistent challenges for food safety and public health (Imade et al., 2021). One effective approach to mitigate this risk is by promoting foods with minimal chemical preservatives, ideally enriched with natural components, which maintain sensory quality while ensuring safety (Soltani et al., 2021). When additive-free options are unavailable, consumers generally prefer products with naturally derived additives over synthetic ones (Perito et al., 2020). In this context, lactic acid bacteria (LAB) emerge as a promising solution, thanks to their ability to produce bioactive compounds that enhance food preservation and safety (Wang et al., 2021). These metabolic capabilities contribute to improved nutritional value, reduced toxin levels, extended shelf life, and enhanced flavour in fermented products (Zheng et al., 2020). In addition to nutrient competition with pathogens, LAB can adhere to host receptors and secrete bacteriocins that target microbial membranes, inhibiting pathogen colonization (Tsouklidis et al., 2020). Beyond live bacteria and their bacteriocins, the concept of postbiotics—non-viable microbial components or metabolic products has gained attention for their health-promoting activities (Hossain et al., 2021). Postbiotics may possess antimicrobial, antioxidant, anti-inflammatory, immunomodulatory, and anti-proliferative effects, holding potential for therapeutic applications (Medema et al., 2015). In food and feed systems, these molecules can suppress pathogen growth and interact with host or microbial communities to help prevent infections (Albarano et al., 2020). The rise of antibiotic-resistant pathogens, particularly methicillin-resistant Staphylococcus aureus (MRSA), poses a significant threat to global public health. MRSA exhibits resistance to β-lactam antibiotics and has a strong ability to form biofilms on various surfaces, contributing to persistent infections and increased mortality (Ali Alghamdi et al., 2023; Silva et al., 2021). In recognition of its severity, the World Health Organization (WHO) classified MRSA as a high-priority pathogen in 2017 due to its association with over 150,000 hospital-acquired infections annually, resulting in significant economic and health burdens. Consequently, there is a pressing need to identify natural antimicrobial alternatives. Recent studies suggest that lactic acid bacteria (LAB) and their metabolites may inhibit biofilm-forming MRSA strains effectively (Jalalifar et al., 2022). To address such concerns, whole genome sequencing (WGS) has emerged as a robust method for evaluating the safety and functionality of probiotic strains. WGS provides insights into antimicrobial resistance genes, virulence determinants, and biosynthetic gene clusters (BGCs) associated with antimicrobial peptide production (Chokesajjawatee et al., 2020). This genomic data not only ensures safety assessment but also allows for identification of strain-specific traits beneficial for therapeutic and food applications (Tenea & Ascanta, 2022). Traditional fermented foods and beverages, especially those produced through indigenous practices, serve as reservoirs of functionally diverse LAB strains. For example, Atingba, a rice-based fermented beverage widely consumed by the Meitei and Kabui communities in Manipur, India, is known for its medicinal and antimicrobial properties. It is prepared using Hamei, a native starter culture rich in LAB such as *Lactobacillus* and *Bifidobacterium*, and has been associated with beneficial effects on human health (Wahengbam et al., 2020; Giri et al., 2018). Among these beneficial microbes, *Lactiplantibacillus plantarum* stands out due to its metabolic versatility and safety profile. Recognized as Generally Recognized as Safe (GRAS) by the U.S. FDA and approved by the European Food Safety Authority, Lp. plantarum is frequently isolated from fermented foods and beverages (Yilmaz et al., 2022). It produces a diverse array of antimicrobial peptides including bacteriocins, lanthipeptides, lasso peptides, and ribosomally synthesized and post-translationally modified peptides (RiPPs), enhancing its therapeutic potential (Huang et al., 2021; Montalbán-López et al., 2021). Furthermore, many strains have demonstrated both narrow- and broad-spectrum antimicrobial activity against various foodborne and clinical pathogens (Goel et al., 2020). These attributes make Lp. plantarum a promising candidate for next-generation probiotics and functional food formulations (Kandasamy et al., 2022). These peptides vary in mode of action, molecular weight, genetic origins, and biochemical properties, allowing different strains to target specific pathogens (Tang et al., 2023). Their membrane-disruptive abilities make them useful agents against sensitive bacteria (Tang et al., 2023). Recent research supports the effectiveness of LAB-secreted metabolites. For example, cell-free supernatants (CFS) from native LAB strains markedly inhibited foodborne pathogens (Tenea et al., 2021). These inhibitory effects stem from direct antimicrobial interactions that impair pathogen growth and colonization. Additionally, the metabolites disrupted cell integrity in target organisms, causing leakage of cytoplasmic aromatic compounds and structural damage to cell membranes and cell walls (Tenea et al., 2020). In this study, we employed WGS to characterize *L. plantarum* C6, with the aim of elucidating genomic features associated with metabolic pathways, probiotic potential, biosynthetic gene clusters (BGCs) responsible for antimicrobial peptide production, and safety-relevant properties. Comparative genomic and phylogenetic analyses were performed to classify *L. plantarum* C6 in relation to other *L. plantarum* strains. Genomic mining enabled the identification of putative antimicrobial compounds and key probiotic determinants implicated in host interaction and competitive exclusion of pathogens. In particular, biosynthetic gene clusters encoding ribosomally synthesized and post-translationally modified peptides (RiPPs) and other bacteriocins were annotated. To validate the genomic predictions, molecular docking analyses were conducted to assess the interaction of the predicted antimicrobial peptides with methicillin-resistant *Staphylococcus aureus* (MRSA) targets, supporting their potential inhibitory activity. The integration of genomic, bioinformatics, and molecular modelling approaches in this study lays a strong foundation for the development of genome-informed, naturally derived antimicrobial agents and probiotic candidates for combating biofilm formation pathogens.

## 2. Materials and methods

### 2.1 Bacterial Strain Lp. plantarum C6 genome collection

The *Lp. plantarum* C6 strain, featuring a 2.78-MB de novo assembled whole-genome sequence, was obtained from the NCBI genome database, GenBank (accession number <u>JBDPMP000000000.1</u>).

### 2.2 Taxonomic profiling and phylogenetic assessment

To determine the genome-based taxonomic placement of the isolate, its genomic sequence was submitted to the Type (Strain) Genome Server (TYGS) (https://tygs.dsmz.de/), which enables genome-wide phylogenetic reconstruction. Species and subspecies identification, as well as the closest type strains, were inferred based on digital DNA–DNA hybridization (dDDH) values, as outlined by Meier-Kolthoff and Göker (2019). Phylogenetic trees were generated using FastME 2.1.6.1, utilizing genome-based distance metrics calculated through the Genome BLAST Distance Phylogeny (GBDP) method (Lefort et al., 2015). Additionally, the Average Nucleotide Identity (ANI) was computed to assess genomic similarity with closely related strains. These ANI values were calculated using the FastANI tool (v1.34), which performs rapid, alignment-free comparisons against reference genomes (Jain et al., 2018).

### 2.3 Genomic analysis including sequencing, assembly, and gene annotation

Whole-genome sequencing (WGS) of *Lactiplantibacillus plantarum* C6 was performed using the Illumina NovaSeq 6000 platform with a paired-end read configuration of 2 × 150 bp. The raw sequence data quality was evaluated using FASTQC (v0.12.0) (Andrews, 2010). SPAdes (v3.15.1) was used for de novo genome assembly (Bankevich et al., 2012). Genome annotation and circular visualization were conducted using the Proksee web server (Grant et al., 2023). Further gene annotation was performed through the NCBI Prokaryotic Genome Annotation Pipeline (PGAP) and the RAST server, which facilitates subsystem-based functional categorization (Tatusova et al., 2016; Aziz et al., 2008). The final circular genome representation was created using CGView available through Proksee (http://proksee.ca/) (Grant et al., 2023). Functional annotation of the predicted proteins was performed using BLASTp (v2.13.0+) to identify sequence similarities against multiple databases including NR, Pfam, UniProt, and COG (Cantalapiedra et al., 2021). Proteins were classified into Clusters of Orthologous Groups (COGs) based on these comparisons. Additionally, KEGG Orthology (KO) assignments and pathway mapping were carried out using the KAAS web server (v2.1) (Kanehisa et al., 2016). Carbohydrate-active enzymes (CAZymes) were identified via the dbCAN2 Meta server, which utilizes the CAZy database for classification (Lombard et al., 2014; Zhang et al., 2018). Orthologous protein cluster analysis was conducted using OrthoVenn2, enabling comparison with reference genomes to detect conserved gene families (Xu et al., 2019).

### 2.4 Investigation of biosafety features in *Lp. plantarum* C6

Antimicrobial resistance (AMR) genes in the *Lactiplantibacillus plantarum* C6 genome were identified using the Resistance Gene Identifier (RGI) tool from the Comprehensive Antibiotic Resistance Database (CARD), applying default parameters with a focus on “Perfect” and “Strict” hits along with high-quality coverage (Alcock et al., 2023). For the detection of CRISPR arrays, associated Cas genes, and prophage elements, analyses were performed using CRISPRCasFinder and the PHASTER tool (Couvin et al., 2018; Arndt et al., 2016). The resulting genome features were visualized in a circular genome map created with Proksee (Grant et al., 2023).

### 2.5 Genome mining of biosynthetic gene clusters (RiPP) and secondary metabolite potential in the C6 genome

To identify genes involved in the biosynthesis of secondary metabolites, the contig FASTA sequences of the *Lactiplantibacillus plantarum* C6 genome were analyzed using BAGEL4 (http://bagel4.molgenrug.nl) for putative bacteriocin gene clusters (van Heel et al., 2018). Additionally, antiSMASH (https://antismash.secondarymetabolites.org/) was employed to screen for biosynthetic gene clusters (BGCs) related to various classes of secondary metabolites (Medema et al., 2011). The contig file was submitted with default parameters, including the KnownClusterBlast feature, which compares identified clusters against the MIBiG database. This enabled prediction of metabolite classes such as nonribosomal peptides (NRPS), polyketides (PKs), and RiPP-like peptides, offering insight into the strain’s antimicrobial biosynthetic potential (Terlouw et al., 2023). Visualization of gene operons was carried out using DNA Features Viewer (https://github.com/Edinburgh-Genome-Foundry/DnaFeaturesViewer), based on the results obtained from BAGEL4 analysis. To detect biosynthetic gene clusters (BGCs) associated with ribosomal synthesized and post-translationally modified peptides (RiPPs), the RiPPMiner-Genome tool was employed (Agrawal et al., 2017). Tanimoto similarity scoring was used to assess structural similarity between molecules, with scores ranging from 0.0–0.2 (indicating minimal similarity) to 0.8–1.0 (representing highly similar or nearly identical chemical structures).

### 2.6 Molecular docking of metabolites with Poly-β-1,6-N-acetyl-D-glucosamine synthase enzyme

A comprehensive in silico approach was employed in July 2025 to investigate the potential anti-MRSA (Methicillin-resistant Staphylococcus aureus) activity of natural small biomolecules derived from the cell-free supernatants (CFS) of *Lactobacillus plantarum*. The study was initiated with literature mining using specific keywords such as “*Lactobacillus plantarum* and MRSA” and “cell-free supernatants, MRSA, and antibiofilm” to identify relevant biomolecules and their reported activities. Databases including PubMed, Scopus, and Google Scholar were used to extract information on bioactive metabolites secreted by *L. plantarum*. For molecular docking studies, the Poly-β-1,6-N-acetyl-D-glucosamine synthase enzyme—an important target involved in biofilm formation in MRSA—was selected. The 3D structure of the enzyme was retrieved from the AlphaFold Protein Structure Database using the ID: *AF-Q6G608-F1-v4*. Prior to docking, active site prediction of the enzyme was carried out using PrankWeb, a web-based tool for ligand-binding site prediction. This helped define the probable binding pocket for docking simulation. Molecular docking was performed using AutoDock Vina integrated within the PyRx 0.9.8 virtual screening environment. Bioactive compounds identified from *L. plantarum* CFS through literature mining were used as ligands. The grid box for docking simulations was configured based on the predicted binding pocket with the following grid parameters:

- **Center Coordinates** – X: 4.7046, Y: 4.1494, Z: −7.2023
- **Dimensions (Å)** – X: 42.9584, Y: 63.2595, Z: 57.8623

All ligand molecules were energy-minimized and converted to the appropriate format prior to docking. The docking results were evaluated based on binding affinity scores and molecular interactions. Finally, ligand-receptor interactions were analyzed and visualized using Discovery Studio Visualizer (2021 edition). Key interactions, including hydrogen bonds, hydrophobic contacts, and pi-stacking, were examined to determine the binding stability and specificity of the biomolecules within the active site of the target enzyme.

## Results and discussion

### 3.1 Taxonomic classification and phylogenetic analysis

The whole genome taxonomy of the Lp. plantarum C6 strain was compared with the closest types of 20 strain genomes available in the TYGS database based on the phylogenomic analysis using genome–genome taxonomy, which revealed that the Lp. plantarum C6 strain showed similarity with other strains such as Lp. plantarum DSM 20174 and Lp. plantarum ATCC 14917, while Lp. brownii WILCCON 0030 and Lactobacillus arizonensis DSM 13273 also showed a close distance (Figure 1). Therefore, the results of the TYGS analysis confirmed the identification of the genes. In this study, the whole genome sequence analysis of Lp. plantarum C6 strain isolated from dairy cheese, a dairy product, revealed a genome size of 3,226,489 bp and a GC ratio of 44.5%. The genome size and GC ratio of Lp. plantarum C6 showed similarity with the previously reported Lp. Plantarum strains isolated from Atingba, a traditional fermented rice-based beverage from Manipur, India. The draft genome comprises 13 contigs, with a total genome size of 3,320,817 bp and a GC content of 44.6%, which aligns well with the typical genomic features of *L. plantarum* species (Huidrom et al., 2024). The relatively larger genome size and higher GC content (guanine– cytosine ratio) observed in *Lactobacillus strains*, including *Lactiplantibacillus plantarum* BRD3A, indicate a greater genetic capacity for environmental adaptability. This means the strain likely harbors a wide range of genes that help it survive under diverse environmental conditions, such as changes in nutrient availability, stress factors, or host interactions (Huidrom et al., 2024; Li et al., 2023). Using the Type Strain Genome Server (TYGS), a genome-based taxonomic analysis was conducted, which revealed that C6 is closely related to known strains such as *L. plantarum* DSM 20174 and *L. plantarum* ATCC 14917. The ANI-based taxonomy was analyzed based on *FastANI* 1.34 values, which revealed that the strain C6 had 95.69-99.39 % genome sequence similarities with the closely related species shown in Figure 2 with a high percentage of similarity with *Lp. plantarum* NMGL2 (99.37%), Lp. plantarum DMDL 9010 (99.39%). The strains *Lp. plantarum* ATCC 14917 (99.22%) followed by Lp. plantarum DSM 20174 (95.69%), *Lp. plantarum* SRCM100442 (98.98%), Lp. plantarum UNQLp11 (98.66%) and Lp. plantarum DMC-S1 (99.37%) belongs to *Lp. plantarum* species, indicating that the C6 strain is from *Lp. plantarum* species. Further confirmation came from FastANI (Average Nucleotide Identity) analysis, a high-resolution method for comparing genome similarity. C6 exhibited 99% nucleotide identity with well-characterized strains *Lp. Plantarum* NMGL2, *Lp. plantarum* DMDL 9010, and *Lp. plantarum* ATCC 14917 well above the species-level threshold, thus, definitively placing it within the *L. plantarum* species. In summary, both phylogenomic analysis and ANI values strongly support that C6 is a strain of *L. plantarum*, and its genomic features point toward enhanced adaptability and functional versatility, making it a promising candidate for industrial or probiotic applications.

**Figure 1.**
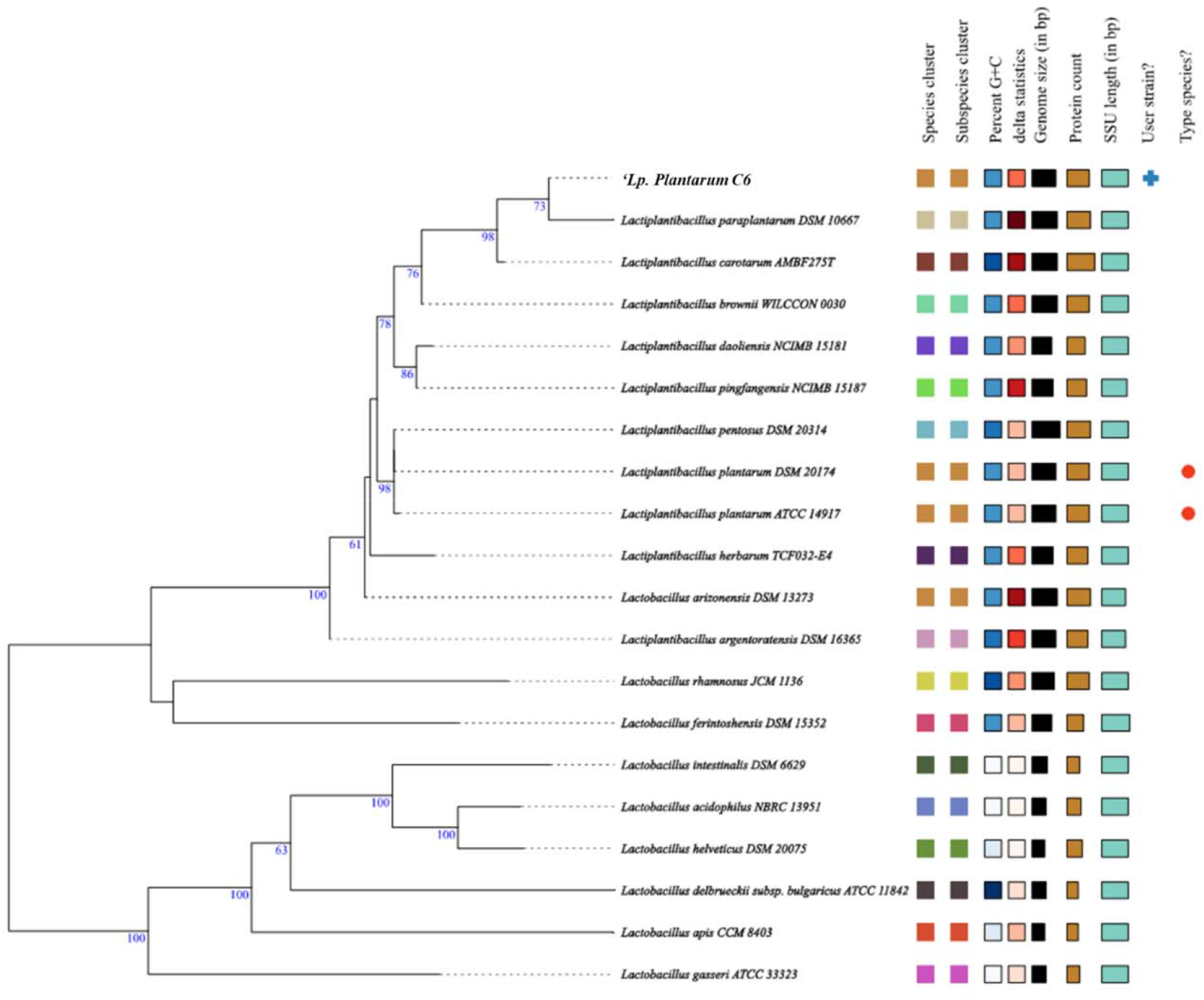
Phylogenetic comparisons of Lactiplantibacillus plantarum C6 with representative genomes of other reference strains carried out in the TYGS webserver indicate. The phylogenetic tree was constructed to determine the evolutionary relationship of *L. plantarum* C6 with closely related *Lactiplantibacillus* and *Lactobacillus* strains based on whole-genome sequence data. Bootstrap values (in blue) indicate branch support. *L. plantarum* C6 clusters closely with *L. plantarum* DSM 20174 and *L. plantarum* ATCC 14917, indicating strong phylogenetic relatedness and high genomic similarity. The icons on the right provide comparative genomic attributes that support this taxonomic placement.

**Figure 2.**
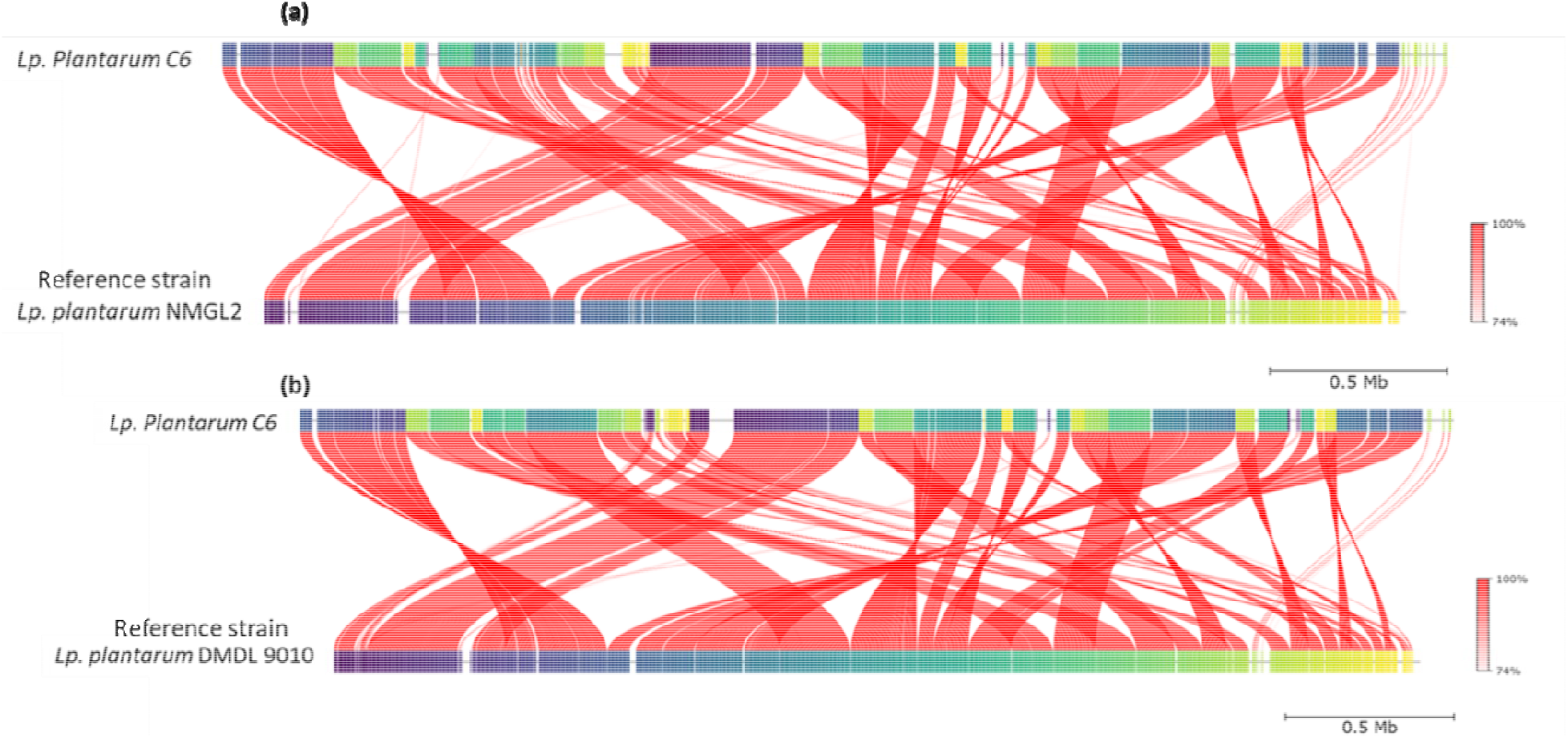
Image of FastANI of Lp. plantarum C6 compared with closely related Lp. plantarum strains that show 99% similarity with *Lp. Plantarum* NMGL2, *Lp. plantarum* DMDL 9010, *Lp. plantarum* ATCC 14917 with reference strains and.

### 3.2 Genome assembly and features

The whole genome sequencing of *Lp. plantarum C6* strain isolated from Indian cheese was performed in the NovaSeq 6000 platform Illumina sequencing. The genome consists of total number of Scaffolds 57 and has an N50 value of 197,852 with a genome size of 3,226,489 bp and a guanine–cytosine (GC) ratio of 44.5%. Based on the Prodigal software for the genomic annotation, a total of 3,027 genes, including total 3,076 protein coding sequence (CDS), out of 30,76 CDS 1234 hypothetical protein, 59 tRNAs, 10 rRNAs, 1 tmRNA, and 2 CRISPR array were predicted. The circular genomic map of *Lp. plantarum C6* was annotated and visualized by Proskee, which showed the characteristics of the genome such as the distribution of the genes on the forward and antisense strands, functional classification of genes, and GC content, as shown in Figure 3.

**Figure 3.**
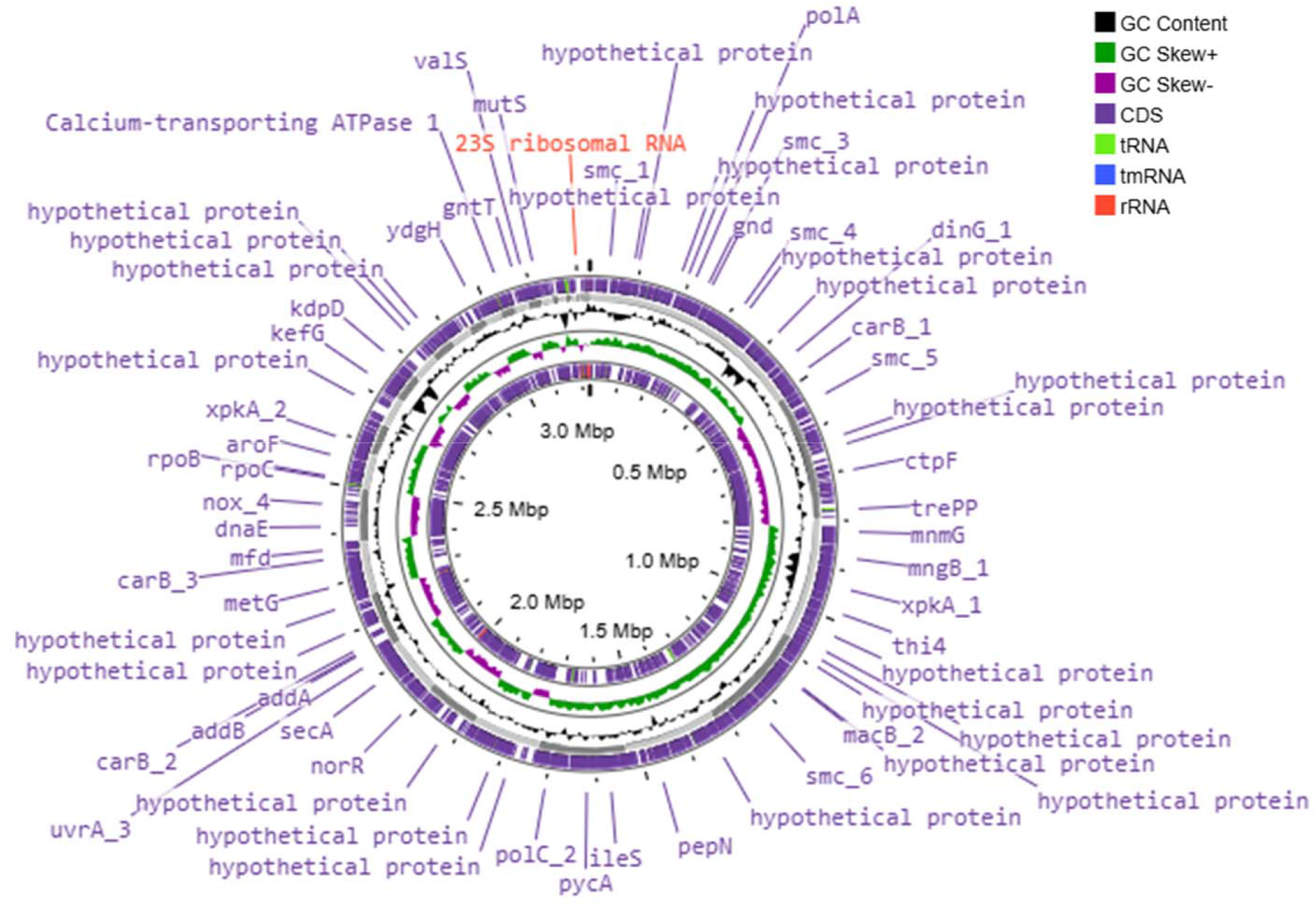
Circular genomic map of *Lactiplantibacillus plantarum* C6 using Proskee View server showing the characteristics features from outside to inner of the circle: CDS, RNA, GC skewness, and GC content are indicated in color code.

The whole-genome sequencing and assembly of *Lactiplantibacillus plantarum* PA21 yielded a draft genome comprising three contigs, with a total genome size of 3,218,706 bp and a GC content of 44.62%, consistent with typical *L. plantarum* strains. The assembly quality was supported by a high N50 value of 1,157,435 bp, indicating substantial contiguity in the genomic structure. Annotation revealed 3,118 protein-coding sequences (CDSs), along with 69 tRNA genes and 14 rRNA genes, signifying a complete set of genes essential for translation and core cellular processes. Additionally, 63 repeat regions were identified, which may play roles in genome plasticity and adaptation (Isaac et al., 2024). These genomic features provide valuable insights into the genetic makeup of C6 and lay the foundation for future functional and comparative genomic studies to explore its probiotic and antimicrobial potential.

### 3.3. Functional characterization and annotation

Simultaneously, all predicted genes from the *Lactiplantibacillus plantarum* C6 genome were subjected to similarity searches against the Pfam, UniProt, and COG databases using BLASTP with an e-value threshold of 1e−5. The results of the annotation are summarized in the table below. Out of a total of 3,027 predicted proteins, 1,869 had significant matches in the Pfam database, 2,168 in Uniprot, and 2,473 in the COG database. Conversely, 1,158 proteins did not match any Pfam entries, 859 showed no hits in UniProt, and 554 lacked matches in the COG database. The functional classification based on COG categories is presented in Table 1. Among the annotated proteins, the highest representation was found in the carbohydrate metabolism and transport category (10.7%), followed by amino acid transport and metabolism (7.8%), translation, ribosomal structure, and biogenesis (7.8%), transcription and cell cycle control (10.5%), cell wall/membrane/envelope biogenesis (6.5%), nucleotide transport and metabolism (4.0%), and coenzyme transport and metabolism (4.9%). These findings highlight the strain’s genetic potential for nutrient metabolism, biosynthesis, and cellular regulation, suggesting its adaptability and functional relevance in diverse environments. In a related study, comprehensive functional annotation of the *Lactiplantibacillus plantarum* BRD3A genome using RAST, COG, and KEGG databases revealed a broad and diverse genetic repertoire. RAST analysis identified 1,985 coding sequences (CDSs) distributed across 339 SEED subsystems, indicating the presence of genes involved in essential metabolic and regulatory functions. COG annotation further classified 2,326 genes into 18 functional categories, with the largest proportion falling under “function unknown”. This significant fraction of uncharacterized genes highlights the genomic novelty of BRD3A and suggests the existence of unique or strain-specific functions that remain to be functionally elucidated (Huidrom et al., 2024).

**Table 1.** Biological subsystem distribution of the genes by RAST annotation for the genome *Lp. plantarum* C6.

| Description | Value | Percent |
| --- | --- | --- |
| Energy production and conversion | 112 | 4.0 |
| Cell cycle control, cell division, chromosome partitioning | 48 | 1.7 |
| Amino acid transport and metabolism | 218 | 7.8 |
| Nucleotide transport and metabolism | 111 | 4.0 |
| Carbohydrate transport and metabolism | 298 | 10.7 |
| Coenzyme transport and metabolism | 138 | 4.9 |
| Lipid transport and metabolism | 104 | 3.7 |
| Translation, ribosomal structure and biogenesis | 218 | 7.8 |
| Transcription | 290 | 10.5 |
| Replication, recombinant and repair | 120 | 4.3 |
| Cell wall membrane envelop biogenesis | 182 | 6.5 |
| Cell motility | 19 | 0.6 |
| Post translation modification, protein turnover, chaperons | 105 | 3.8 |
| Inorganic ion transport and metabolism | 128 | 4.6 |
| Secondary metabolites biosynthesis, transport and catabolism | 24 | 0.8 |
| General function prediction only | 239 | 8.6 |
| Function unknown | 124 | 4.4 |
| Signal transduction mechanism | 139 | 5.0 |
| Intracellular trafficking, secretion, and vesicular transport | 22 | 0.7 |
| Defense mechanism | 74 | 2.6 |
| Extracellular structure | 4 | 0.1 |
| Mobilome: prophages, transposons | 41 | 1.4 |
| Cytoskeleton | 3 | 0.1 |

The functional genes are classified into several categories for energy production and conversion (C: 112), cell cycle control, cell division, chromosome partitioning (D: 48), amino acid transport and metabolism (E: 218), nucleotide transport and metabolism (F: 111), carbohydrate transport and metabolism (G: 298), coenzyme transport and metabolism (H: 138), lipid transport and metabolism (I: 104), translation, ribosomal structure and biogenesis (J: 218), transcription (K: 290), replication, recombinant and repair (L: 120), cell wall membrane envelop biogenesis (M: 182), cell motility (N: 19), post translation modification, protein turnover, chaperons (O: 105), inorganic ion transport and metabolism (P: 128), secondary metabolites biosynthesis, transport and catabolism (Q: 24), general function prediction only (R: 239), function unknown (S: 124), signal transduction mechanism (T: 139), intracellular trafficking, secretion, and vesicular transport (U: 22), defense mechanism (V: 74), extracellular structure (W: 4), mobilome: prophages, transposons (X: 41), cytoskeleton (Z: 3). The highest proportion of genes was present under carbohydrate transport and metabolism (G: 298) and transcription (K: 290), while cytoskeleton (Z: 3) was associated with a small proportion of genes shown in Figure 4a. In another study, the core genome analysis of *Lactiplantibacillus plantarum* BRD3A revealed the presence of 2,358 orthologous proteins, representing a set of highly conserved essential genes shared across most *L. plantarum* strains. These core genes are predominantly involved in critical cellular functions such as the cell cycle, carbohydrate metabolism, and protein metabolism, consistent with previous findings (Huang et al., 2020). The conservation of these genes underscores their fundamental role in maintaining cellular homeostasis and metabolic versatility, enabling *L. plantarum* strains to adapt to a wide range of environmental niches. The robust representation of metabolic and regulatory genes in the core genome highlights C6 potential for stable performance in industrial and probiotic applications.

**Figure 4.**
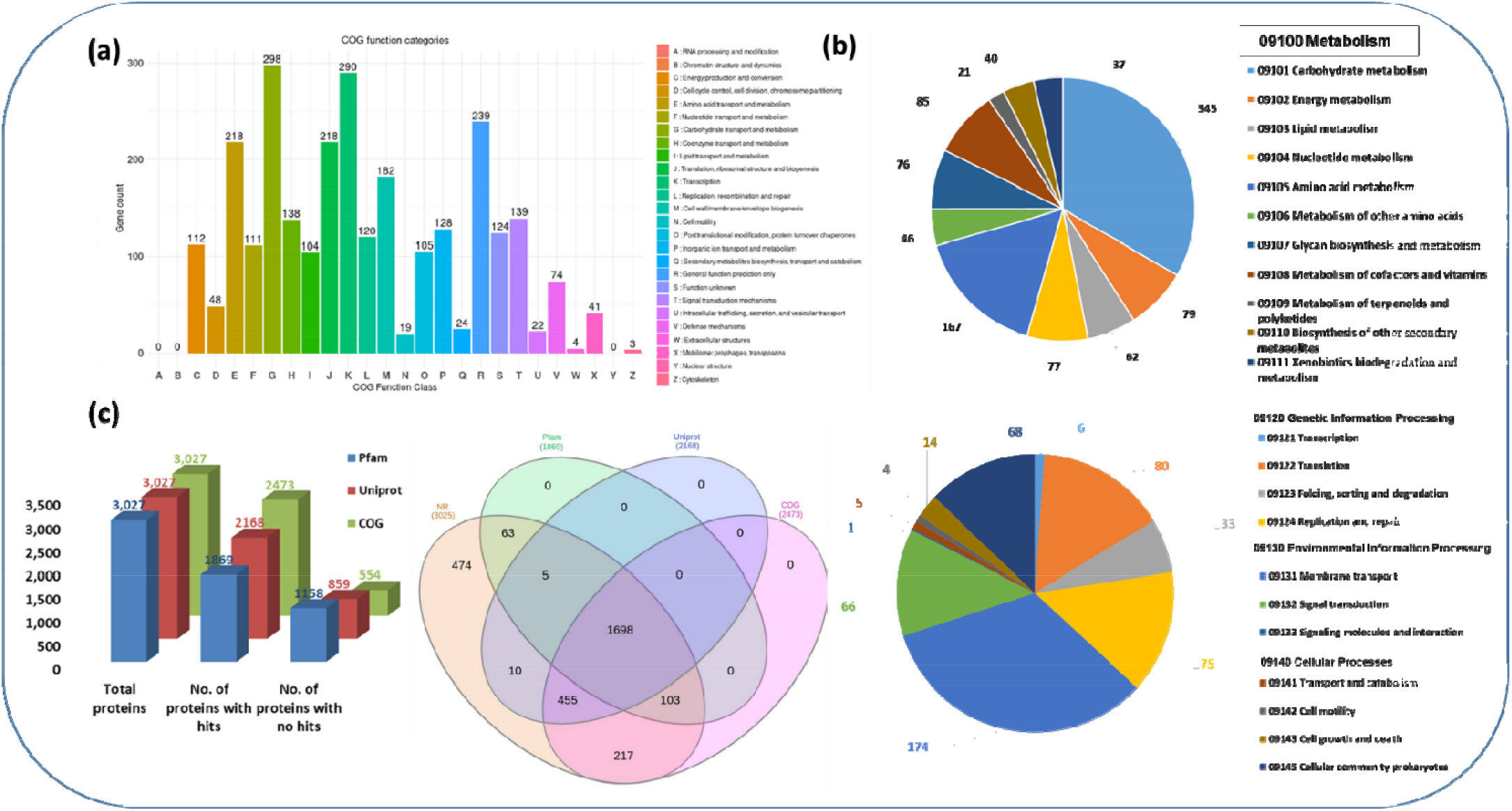
**(a)** Blastp 2.13.0+ prediction of Cluster of Orthologous group (COG) functional categories to the proteins C6 strain, **(b)** KEGG orthology (KO) categories of identified protein-coding genes in the *Lp. plantarum* C6, and **(c)** Comparative analysis of gene annotation in different databases was represented in Venn diagram *Lp. plantarum* C6.

Gene ontology (GO) annotation was obtained for NR database annotated proteins using OmicsBox 3.0. GO sequence distributions helps in specifying all the annotated nodes comprising of GO functional groups. The GO sequence distribution was analysed for all the three GO domains i.e. biological processes, molecular function and cellular components. A total of 1341 genes were assigned GO terms in C6 sample. Gene distribution in different categories, genes annotated in each category such as biological process (914), cellular component (689) and molecular function (1114).

Furthermore, Pathway analysis, ortholog assignment and mapping of genes to the biological pathways were performed using KEGG automatic annotation server (KAAS). All predicted genes sequences from C6 sample were compared against the KEGG database using BLASTP with threshold bit-score value of 60. The mapped proteins represented metabolic pathways of major biomolecules such as carbohydrates, lipids, nucleotides, amino acids, glycans, cofactors, vitamins, terpenoids, polyketides, etc. The proteins also represented the genes involved in metabolism, genetic information processing, and environmental information processing and cellular processes. The distribution of genes assigned to different pathways is shown in Figure 4 b. It suggested that the highest number of genes encoding proteins (345) are responsible for carbohydrate metabolism (KO09101), followed by others for protein families: genetic information process (KO09121-KO09124) (194), environmental information processing membrane transport (09131) (174), signal transduction (09132) (66) and cellular community (KO09145) (68) Figure 4 b. In a similar study, functional categorization of the *L. plantarum* BRD3A genome using BlastKOALA revealed that 1,451 annotated genes were classified into 23 KEGG functional categories. Among these, the largest group (214 genes) was associated with carbohydrate metabolism (KO09101), underscoring the strain’s strong potential for utilization and processing of diverse carbohydrate substrates, a characteristic trait of lactic acid bacteria. This was followed by genes involved in genetic information processing (KO09182) with 201 genes, and those related to signalling and cellular processes (KO09120) (176 genes), suggesting robust regulatory and stress response systems.

Additionally, 140 genes were classified under environmental information processing (KO00130), indicating the strain’s ability to perceive and respond to external environmental stimuli. These annotations reinforce the strain’s capacity to engage in complex biochemical activities and suggest potential applications in functional foods, biotechnology, or antimicrobial development. The combination of conserved core functions and uncharacterized genetic elements points to *L. plantarum* BRD3A as a promising candidate for future functional genomics and probiotic research (Huidrom et al., 2024). These findings are consistent with previous reports indicating that lactic acid bacteria (LAB), including *L. plantarum*, are naturally enriched in genes involved in carbohydrate metabolism (Ghattargi et al., 2018). Such genomic traits not only support efficient energy production but also contribute to the strain’s potential role in fermentation processes, gut colonization, and probiotic functionality. The robust metabolic network encoded by C6 highlights its promise as a functionally active probiotic strain suitable for application in food biotechnology and health-promoting formulations.

Comparative analysis of genes annotation in different databases was represented in Venn diagram. The Venn diagram represents a comparative analysis of gene annotations across four major databases: NR (Non-Redundant Protein Database), Pfam, Uniprot, and COG. Each circle corresponds to proteins annotated by a specific database, and the overlaps indicate the number of genes commonly identified among them. A total of 3025 proteins were analyzed. 1698 proteins were commonly annotated by all four databases, indicating strong consensus in functional prediction. 474 proteins were uniquely identified by NR, and 217 proteins were exclusive to COG, reflecting database-specific annotations. 455 proteins were shared between NR, Uniprot, and COG, while 103 proteins were common between Uniprot and COG only. A smaller number of proteins (63, 10, 5) were shared among two or three databases. Notably, there were no proteins uniquely annotated by Pfam or Uniprot alone, suggesting that these databases may rely on conserved domains or curated entries that overlap significantly with others as show

### 3.4 Presence of probiotic-related genes

The *Lp. plantarum* C6 genome harbors probiotic-related genes responsible for proteins coding genes involved in stress response (temperature, bile, pH) and adhesion genes, which includes cold-shock protein (cspA) that contains three genes related to survival under low temperatures. Heat-shock proteins encode 16 genes, which include molecular chaperones (*dna*K, *dna*J, *hsl*O, *Grp*E, *gros*ES, *htp*X) and protease encoding genes (*Clp*E, *Clp*P, *Clp*X, *Clp*C, *hsl*V, *hslO*). For acid tolerance, there are genes (*atp*C, *atp*D, *atp*G, *atp*A, *atp*H, *atp*F, *atp*E, *atp*B, and *gad*B) encoding resistance in low pH conditions and bile stress, which include three encoding proteins (choloylglycine hydrolase (cbh)) and adhesion-related genes (*gpr, tuf*, dapF and enolase), which indicates the strain’s high adhesion ability, as shown in

Table 2. Studies have shown that *Lactiplantibacillus plantarum* contains genes linked to probiotic functions such as resistance to thermal fluctuations, acidic and bile-rich conditions, and host tissue adherence (Huidrom et al., 2024; Kandasamy et al., 2022). Consistent with these observations, our findings indicate that the C6 strain carries genetic features that may enable it to endure various stress factors, suggesting its potential adaptability within the gut microbiome and gastrointestinal environment.

### 3.5 Prediction of carbohydrate-active enzymes

The genomic analysis of *Lactiplantibacillus plantarum* C6 revealed a comprehensive set of carbohydrate-active enzymes (CAZymes), encompassing 102 genes categorized into five primary CAZy classes, based on annotation through the dbCAN3 platform. These include 58 genes for glycoside hydrolases (GHs), 33 for glycosyltransferases (GTs), 5 for carbohydrate esterases (CEs), 2 for carbohydrate-binding modules (CBMs), and 4 for auxiliary activity (AA) enzymes. The high representation of GH and GT families suggests a robust capability for carbohydrate utilization and interactions with the host environment. Comparable CAZyme gene profiles have been described in other *L. plantarum* strains (Huidrom et al., 2024; Mollova, 2023). These enzymes contribute to the biosynthesis, transformation, and breakdown of complex polysaccharides and glycoconjugates, which are crucial for the physiology of lactic acid bacteria (Kandasamy et al., 2022). Notably, GHs play critical roles in biological functions such as sugar metabolism, energy production, and intracellular signaling pathways (Zhao et al., 2023).

**Table 2.**
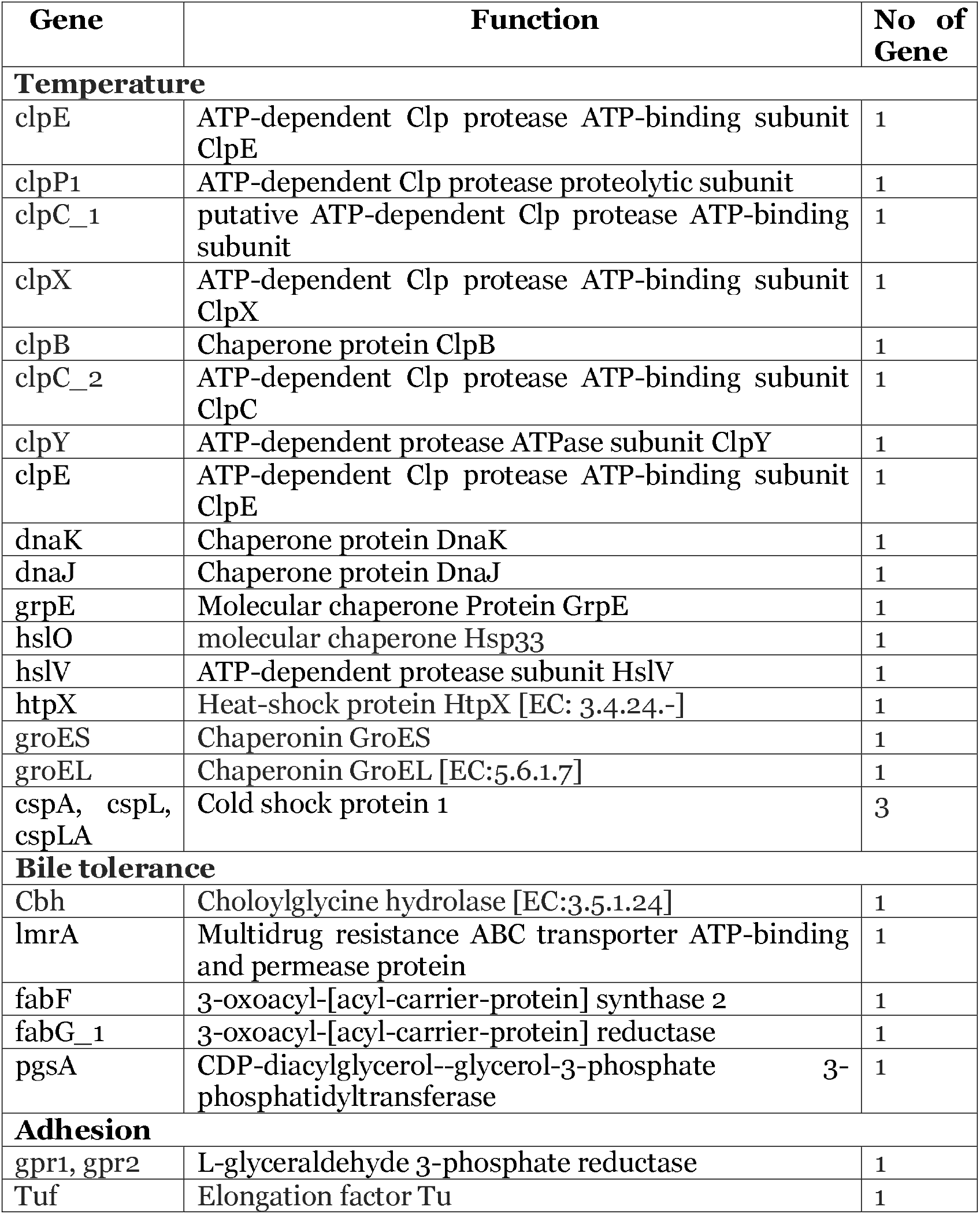
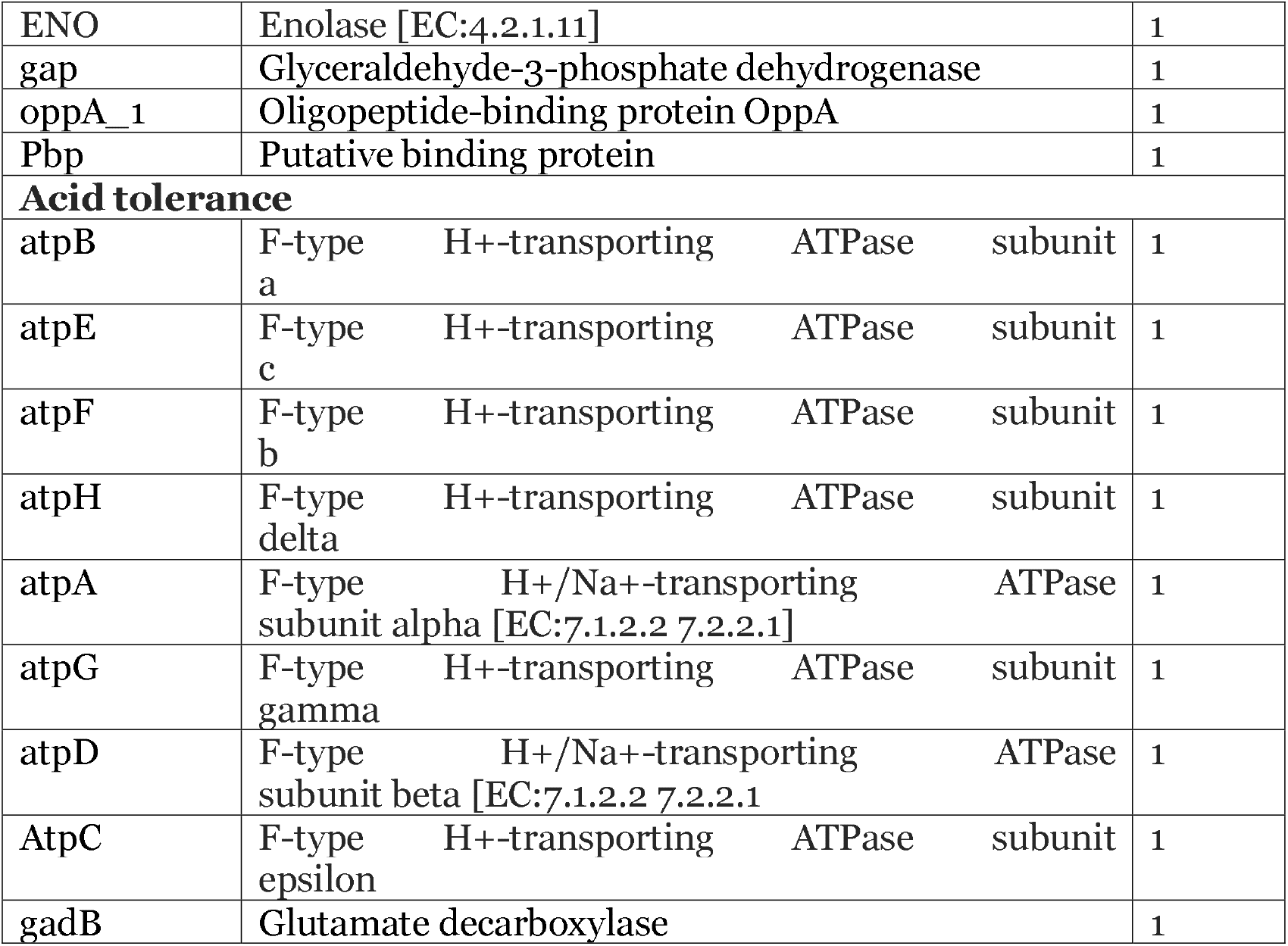
Annotation of probiotic-related genes identified in the *Lp. plantarum* C6 genome.

### 3.6 Safety assessment of the *Lp. Plantarum* C6 strain and prophage region and CRISPR array

The C6 genome predicted from the CARD database revealed two strict hits: (i) *van*H coding gene (glycopeptide antibiotic) with 35.02% identity and (ii) *van*Y, 31.93% identity which gives it small multidrug resistance (SMR), leading to its classification in the drug class as a disinfecting agent and an antiseptic with 31.93% identity of the matching regions (Figure 5 a & Supplementary file 1). Comparable findings were also reported by Huidrom et al. (2024) and Gueimonde et al. (2013). The C6 strain genome of the prophage sequence using PHASTER revealed two prophage regions, comprising two intact (Figure 5 a). The sizes of these regions were 45Kb and 46.7Kb, respectively. The intact prophage (Region 1) and (Region 2) exhibited the protein count of 53, 62 with a GC content of (41.30% and 38.75%) with the most common: PHAGE_Lactob_Sha1_NC_019489 (38), PHAGE_Lactob_phig1e_NC_004305 (9) in the prophage (Region 1 & 2). Comparable findings have been reported in the genomes of other strains, supporting the consistency of these features across L. plantarum species (Huidrom et al., 2024; Zhang et al., 2019). Two intact prophage regions, designated as Region 1 and Region 2, were identified based on the presence of integrated phage elements. These regions harbor key phage-related genes, including integrase, head and tail structural proteins, terminase, portal protein, and holin, suggesting the potential for functional phage activity. The presence of these elements may also contribute to the strain’s ability to resist invasion by foreign prophages (Tenea and Ortega, 2021; Zhang et al., 2019). The purple colour (tRNAs) and attachment sites (blue) are marked in proximity to prophage regions. GC skew and GC content tracks in the innermost rings (green, purple, and black) reveal genomic variation and help localize horizontal gene transfer events, such as prophage insertions. The presence of intact prophage regions suggests that *L. plantarum* C6 may harbor inducible prophages, which could play roles in gene transfer, bacterial fitness, or antimicrobial activity showed in Figure 6. Two CRISPR arrays were identified in the *Lp. Plantarum* C6, one of which is in contig C6 C6_Scaffold_4_1 (start 217,558-end 217, 755 bp) with 198 bp repeat matching a consensus sequence with C6_Scaffold_18_1 with (start 38,702 bp-end 54,770 bp) with 86 bp repeat length evidence level showed in Figure 5 b & Table 3. Consistent findings were observed by Huidrom et al. (2024) in *L. plantarum* strains. The CRISPR/Cas system contributes significantly to bacterial immunity, particularly by limiting horizontal gene transfer events associated with antimicrobial resistance and virulence traits (Rodrigo-Torres et al., 2019). Two CRISPR arrays were identified in the *Lp. Plantarum* C6, one of which is in contig C6 C6_Scaffold_4_1 (start 217,558-end 217, 755 bp) with 198 bp repeat matching a consensus sequence with C6_Scaffold_18_1 with (start 38,702 bp-end 54,770 bp) with 86 bp repeat length evidence level showed in Figure 5 b & Table 3. Consistent findings were observed by Huidrom et al. (2024) in *L. plantarum* strains. The CRISPR/Cas system contributes significantly to bacterial immunity, particularly by limiting horizontal gene transfer events associated with antimicrobial resistance and virulence traits (Rodrigo-Torres et al., 2019). The purple colour (tRNAs) and attachment sites (blue) are marked in proximity to prophage regions. GC skew and GC content tracks in the innermost rings (green, purple, and black) reveal genomic variation and help localize horizontal gene transfer events, such as prophage insertions. The presence of intact prophage regions suggests that *L. plantarum* C6 may harbor inducible prophages, which could play roles in gene transfer, bacterial fitness, or antimicrobial activity showed in Figure 6.

**Table 3.**
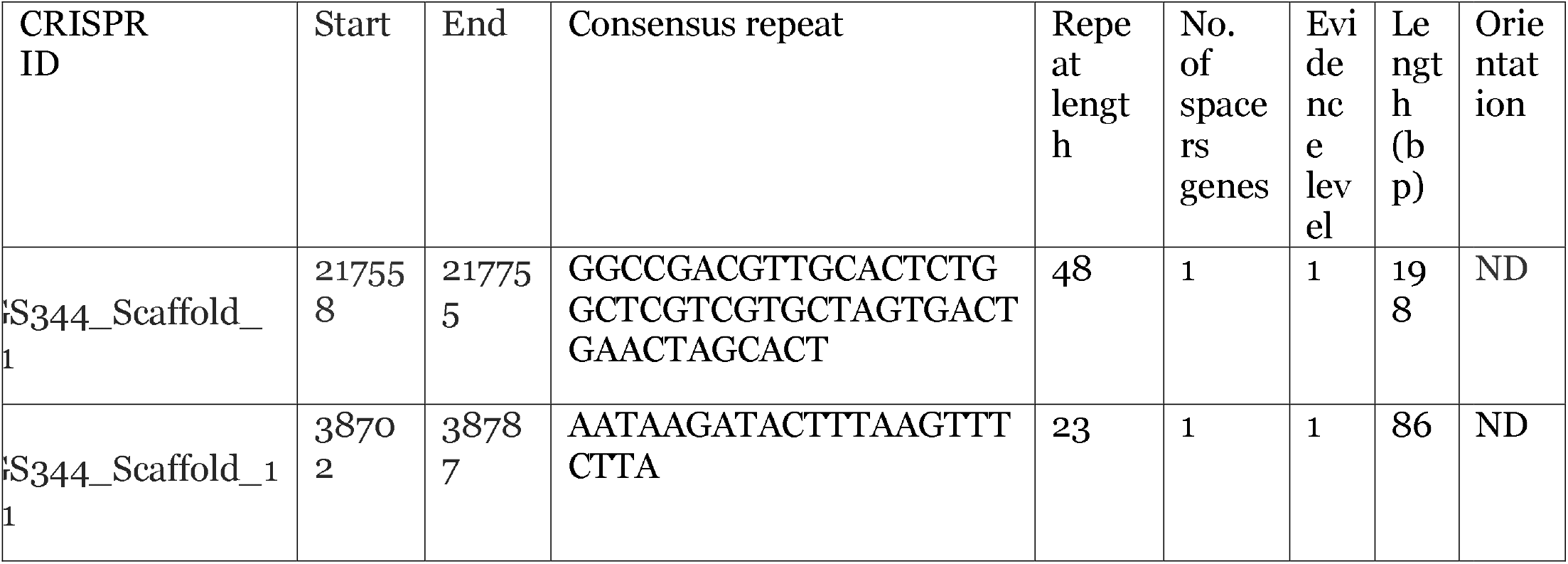
Lp. Plantarum C6 strain CRISPR array system using CRISPRCasFinder.

**Figure 5.**
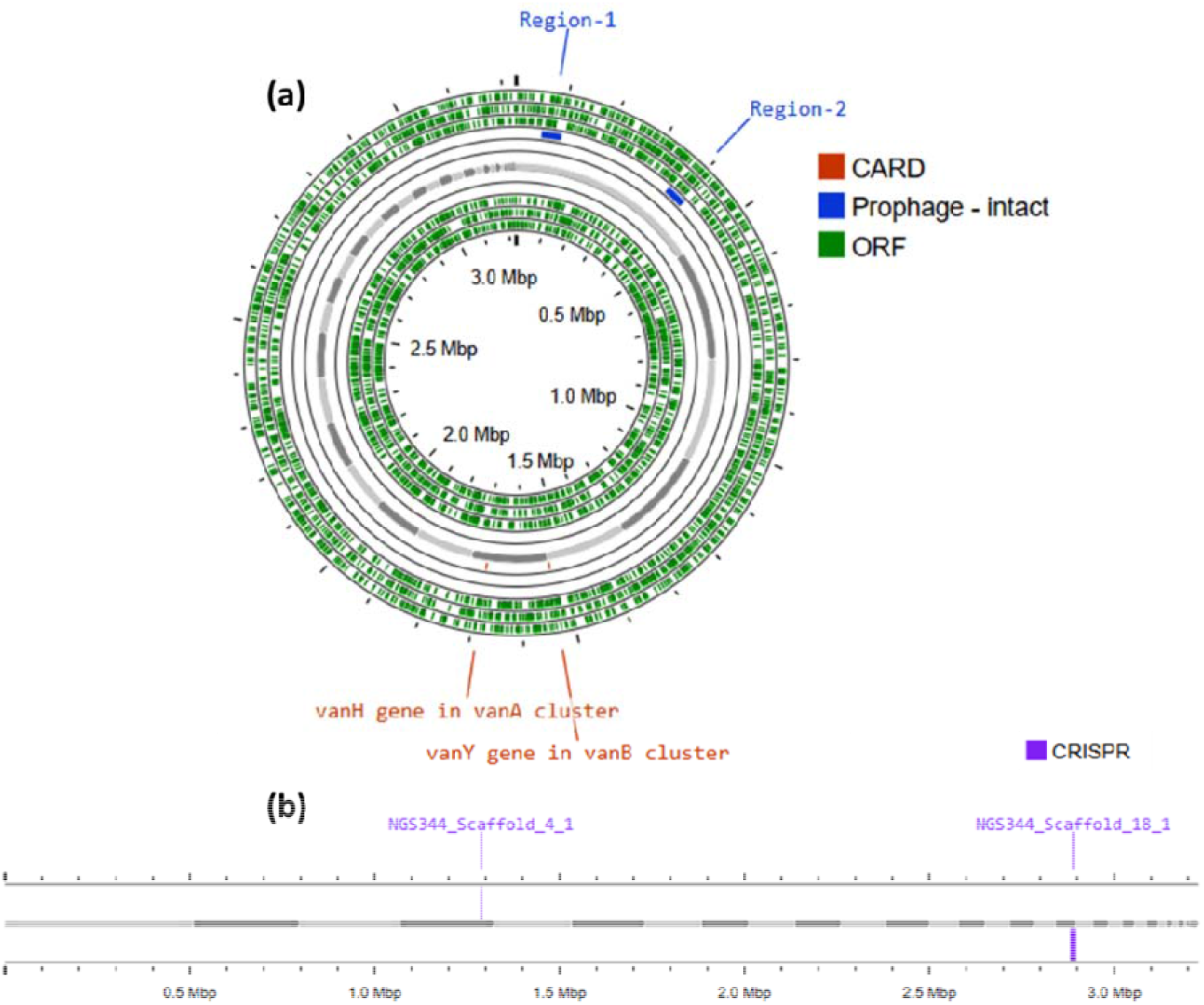
**(a)** Circular genome annotation of *Lactiplantibacillus plantarum* C6 Including Prophages, Resistance Genes, and **(b)** CRISPR Elements

**Figure 6.**
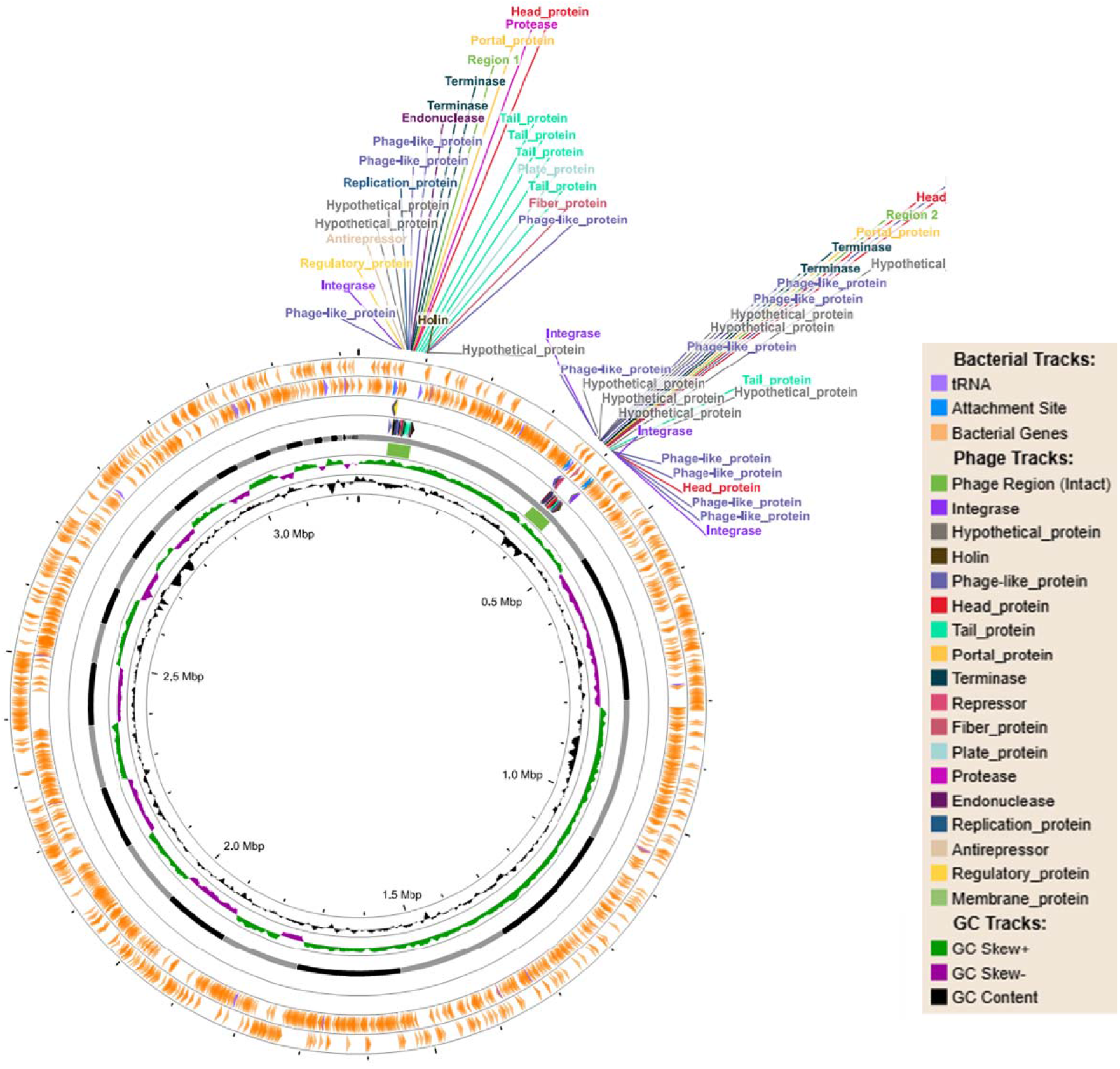
The circular genome map illustrates the organization and annotation of the *L. plantarum* C6 genome with a specific focus on prophage regions and associated genes.

### 3.7. Prediction of biosynthetic gene clusters for bacteriocins

There is ever-growing evidence that bacteriocins are among the bioactive compounds produced by *L. plantarum*, responsible for the inhospitable environment towards MDR bacteria. The biosynthetic gene clusters (BGCs) responsible for bacteriocin production in *Lactiplantibacillus plantarum* C6 were predicted using the BAGEL4 web server, which identified a key area of interest (AOI) located within Scaffold_11.3.AOI_01 and Scaffold_15.5.AOI_01. This area encodes multiple Plantaricin peptides—Plantaricin_E, F, J, K, and N-with high bit scores and significant E-values, confirming their identity (Table 4).

**Table 4.**
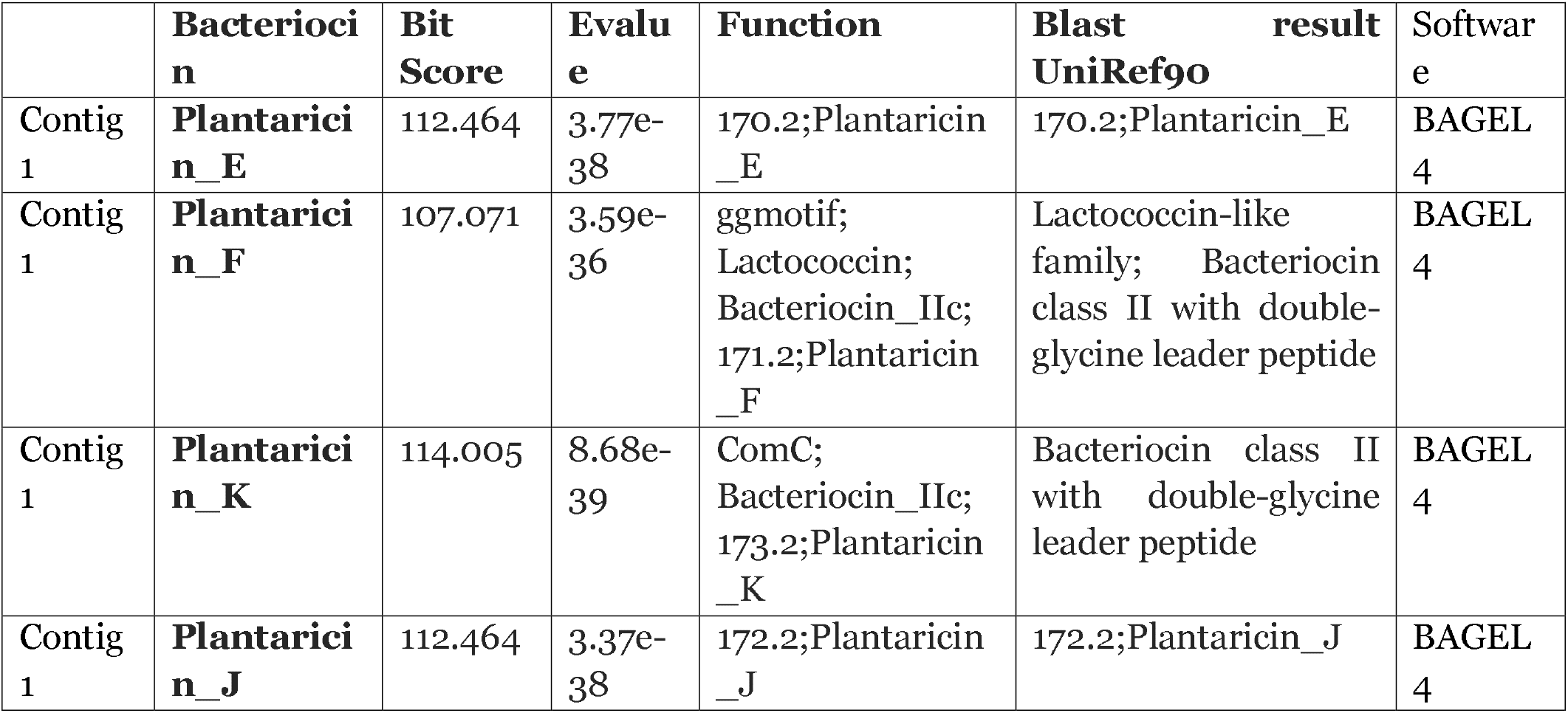
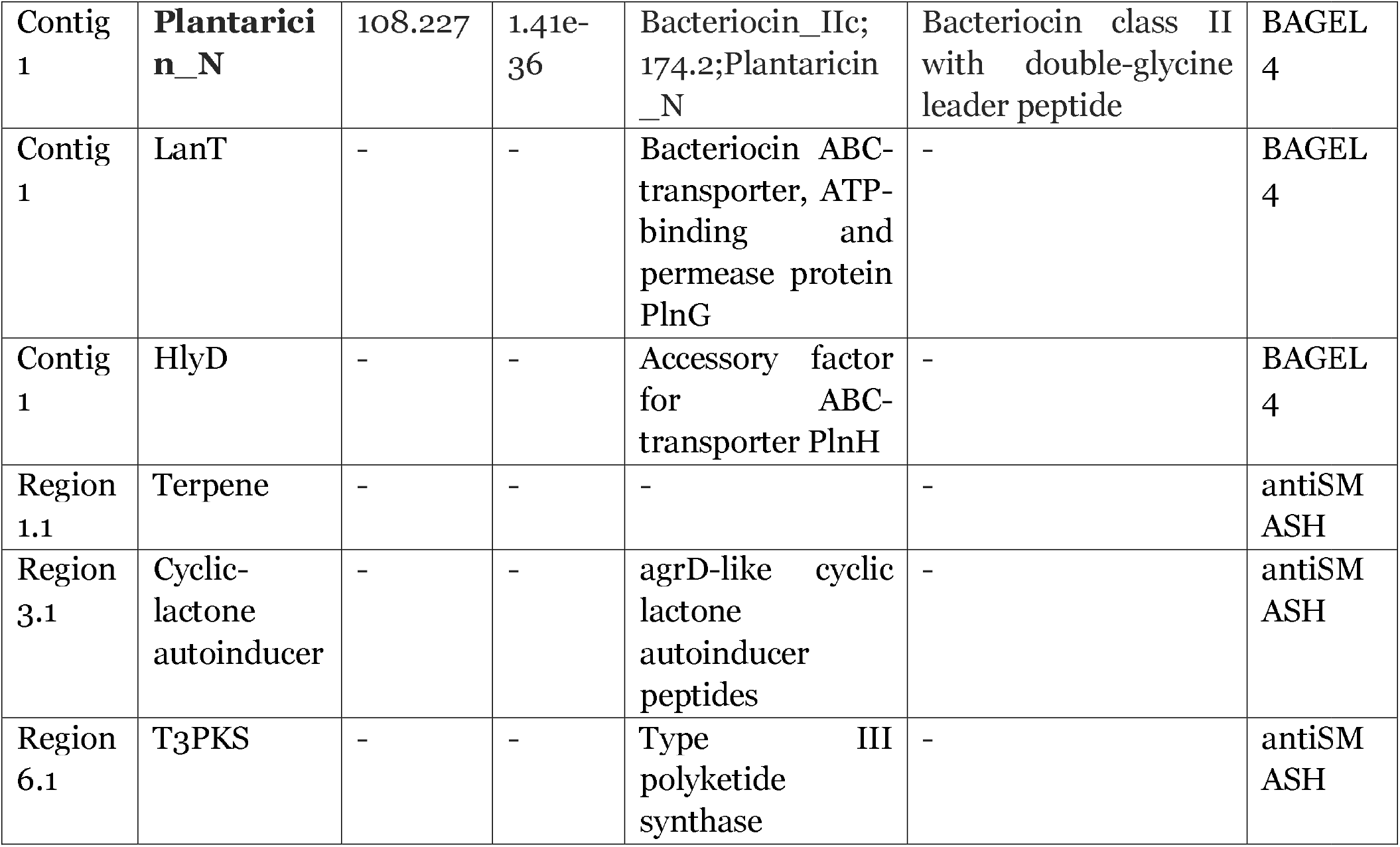
The genome-based identification of L. plantarum C6 biosynthetic gene clusters (BGCs)

These peptides belong to the lactococcin-like family and are characterized by the presence of double-glycine leader sequences and β-chain-like core peptide structures, features typical of class II bacteriocins—small, unmodified, and heat-stable antimicrobial peptides (<10 kDa) (Acedo et al., 2018; Nissen-Meyer et al., 2010). The presence of these gene clusters suggests that *L. plantarum* C6 is a potential class II bacteriocin producer. Similar findings have been documented in previous studies (Goel et al., 2020; Huidrom et al., 2024; Rodrigo-Torres et al., 2019; Tenea and Ortega, 2021), reinforcing the probiotic relevance of such strains. Bacteriocin-producing commensals play a vital role in preventing pathogen colonization and maintaining microbial homeostasis. Moreover, as highlighted by Krauss et al. (2023), understanding the metabolic burden and evolutionary adaptations associated with bacteriocin biosynthesis is essential for optimizing strain fitness and guiding the development of targeted probiotic and microbiome-editing strategies.

The antiSMASH database analysis revealed three active metabolite biosynthetic regions. Although the function of the *plnN* gene was not explored in detail, its localization within the bacteriocin operon suggests a potential regulatory role in bacteriocin production. Furthermore, whole-genome analysis identified biosynthetic gene clusters (BGCs) for secondary metabolites such as terpenes, type III polyketide synthases (T3PKS), and cyclic-lactone inducers—compounds commonly associated with antimicrobial activity. These BGCs have been implicated in the inhibition of indicator strains, as reported in previous studies (Tenorio-Salgado et al., 2021; Hegemann et al., 2018). Collectively, the genome mining results indicate that *Lactiplantibacillus plantarum* PA21 possesses the genetic potential to synthesize diverse antimicrobial compounds showed in Table 4.

Additionally, HlyD bacteriocin is an accessory factor for ABC transporter PlnH (LanT) and Bacteriocin ABC transporter, Bacteriocin ABC-transporter, ATP binding and permease protein PlnG, Immunity protein PlnI, belong to membrane-bound protease CAAX family (Figure 7a). GlyS, marked in blue, is involved in post-translational modification of the core peptide, a hallmark feature of RiPP biosynthesis. Genes in orange and yellow represent functions associated with regulation and leader peptide cleavage, critical steps in peptide activation and export. The additional open reading frames (ORFs) predicted the presence of putative bacteriocins immunity protein found in the same bacteriocin gene clusters PlnC and PlnD, response regulator and activator for transcription factor and bacteriocin production-related histidine kinase within ORFs, as shown in Figure 7b & supplementary file 2.

**Figure 7.**
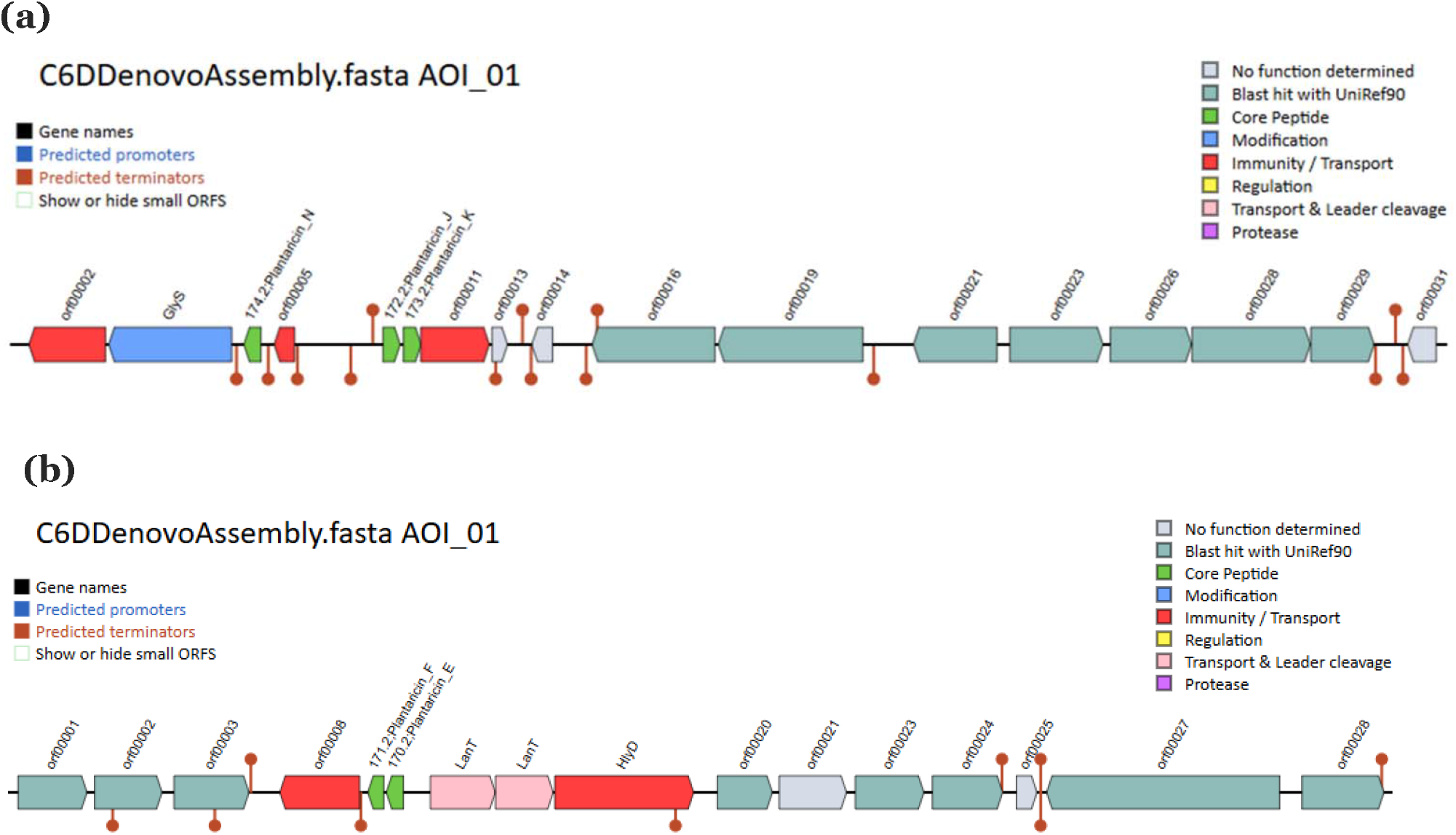
This annotated region represents a predicted biosynthetic gene cluster (BGC) from *Lactiplantibacillus plantarum* C6, (a) possibly encoding a **bacteriocin Plantaricin_K** (Bit Score=114.005) and (b) **Plantaricin_F** (Bit Score=107.071).

### 3.8. *RiPP* Prediction from C6 Genome

Microorganisms produce diverse natural products through dedicated biosynthetic gene clusters (BGCs) groups of co-localized and co-regulated genes that encode the enzymes needed to synthesize these bioactive compounds, such as antibiotics, bacteriocins, and secondary metabolites (Harwood et al., 2018). Biosynthetic gene clusters (BGCs) are mainly classified into two types: ribosomally synthesized and post-translationally modified peptides (RiPPs), produced from precursor peptides modified after translation, and non-ribosomal peptides (NRPs), synthesized independently of the ribosome by large enzyme complexes called non-ribosomal peptide synthetases (NRPSs) (Han et al., 2024). RiPPs (ribosomally synthesized and post-translationally modified peptides), such as lanthipeptides and lasso peptides, are produced through ribosomal synthesis followed by enzymatic modifications, while NRPs (non-ribosomal peptides) are assembled by multi-enzyme complexes without ribosomes. RiPPs are gaining attention as innovative solutions across healthcare and agriculture due to their potential as antibiotics, antivirals, anticancer agents, eco-friendly biopesticides, and natural food preservatives like nisin (Harris et al., 2020).

In our results, genome annotation revealed a RiPP biosynthetic gene cluster located on contig 4 (positions starts at 89,766 and ends at 128,298) of the *L. plantarum* C6 genome. This cluster belongs to the head-to-tail cyclized RiPP class and encodes a Class IId bacteriocin with a cyclical uberolysin-like biosynthetic domain (ORF_11) (Figure 8).

**Figure 8.**
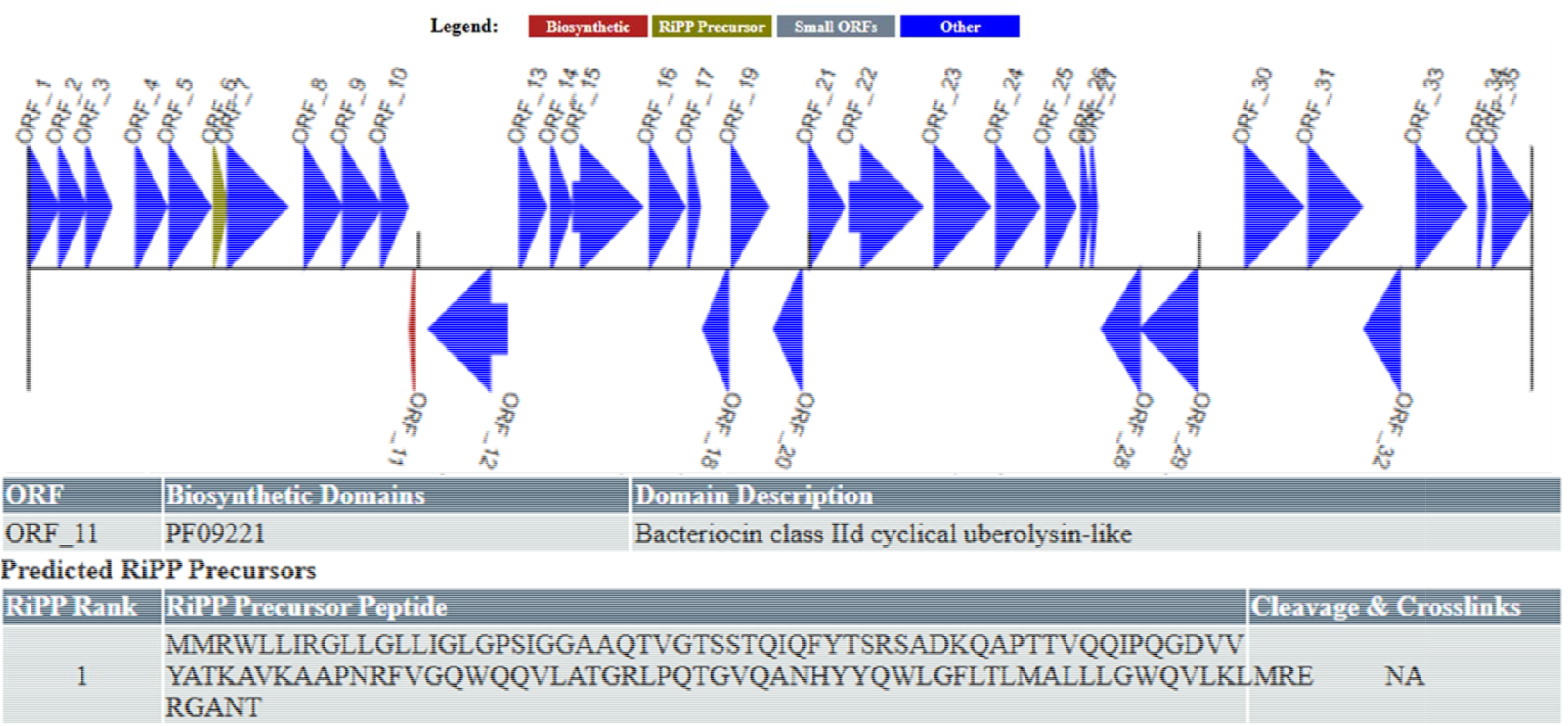
Biosynthetic gene cluster and predicted RiPP domain in the C6 genome. Red arrow: biosynthetic; yellow arrow; RiPP precursor; grey arrow: small ORFs; blue arrow: other small ORFs. ORF: open reading frame.

These bacteriocins are membrane-active peptides commonly produced by Firmicutes and function by disrupting bacterial membranes (Zhang et al., 2022). Notably, lassomycin, a ribosomally synthesized cyclic peptide with strong basicity has demonstrated potent bactericidal activity against both actively growing and dormant *Mycobacterium tuberculosis*, including drug-resistant strains. This highlights its therapeutic potential as a novel antimicrobial agent (Gavrish et al., 2014). However, not all predicted peptides are necessarily expressed or detectable under specific experimental conditions. Although biosynthetic genes may be transcribed, post-transcriptional events such as RNA degradation, editing, or alternative splicing can hinder the translation and eventual production of the predicted metabolites (Piazzi et al., 2023).

RiPPs analysis, a sequence similarity search using the RiPPMiner-Genome web server, identified that the RiPP precursor peptide in the *Lactiplantibacillus plantarum* C6 genome showed homology with three known RiPPs: Butyrivibriocin AR10, Microcyclamide, and Microcyclamide McaE2. Comparative analysis highlighted a diversity of antimicrobial peptides from both bacterial and cyanobacterial origins. Butyrivibriocin AR10, produced by Butyrivibrio fibrisolvens, is a head-to-tail cyclized bacteriocin composed of a 23-amino-acid leader peptide and a 58-amino-acid core region. Its cyclized structure confers enhanced stability and potent antimicrobial activity (as shown in Table 5). In contrast, Microcyclamide and Microcyclamide McaE2 produced by Microcystis aeruginosa NIES-298 belong to the cyanobactin class of RiPPs, which are commonly found in cyanobacteria and exhibit diverse bioactivities, including antimicrobial potential. These RiPPs contain repeated HCATIC motifs, indicatives of heterocyclization into thiazole/oxazole rings, which enhance bioactivity and structural rigidity. Microcyclamide biosynthesis involves the CyaG modification system, while McaE2 carries conserved core motifs at positions 43–48 and 57– 62.

**Table 5.** Sequence Similarity of the C6 Genome RiPP Precursor Peptide with Known RiPPs.

| Subject | Identity | Alignment length | Alignment start | Alignment end | e-value |
| --- | --- | --- | --- | --- | --- |
| Butyrivibriocin AR10 | 42.11 | 19 | 11 | 29 | 4.9 |
| Microcyclamide | 25.00 | 32 | 23 | 42 | 8.9 |
| Microcyclamide McaE2 | 25.00 | 32 | 23 | 42 | 9.5 |

A similar study by Molina et al. (2025) reported the identification of a RiPP biosynthetic gene cluster in contig 8 (positions 20,529 to 61,647) of the *UTNGt2* genome. This cluster belongs to the head-to-tail cyclized RiPP class and encodes a Class IId cyclical uberolysin-like bacteriocin (ORF_11). These bacteriocins are membrane-active peptides commonly produced by members of the *Firmicutes* phylum (Zhang et al., 2022). Previous studies have demonstrated that lassomycin, a highly basic, ribosomally synthesized cyclic peptide, possesses potent bactericidal activity against both actively replicating and dormant *Mycobacterium* species, including multidrug-resistant strains of *M. tuberculosis* (Gavrish et al., 2014). Collectively, these peptides highlight the structural and functional diversity of natural antimicrobials showed in Table 5.

Structural Similarity of RiPP Precursor Peptide: A closest-structure similarity search of the RiPP precursor peptide from the Lactiplantibacillus plantarum C6 genome revealed high structural similarity with several known RiPPs. Notably, the peptide showed a perfect Tanimoto score of 1 with Streptococcin A M49, Streptococcin A FF22, Salivaricin 9, Macedocin, and Ericin S. High similarity was also observed with Lacticin 3147 A1 (Tanimoto score: 0.967742), Subtilin (0.966667), Entianin (0.966667), Nisin Q (0.9375), and Nisin F (0.9375). All structurally similar RiPPs belong to the LanthipeptideA or LanthipeptideB classes, as detailed in Table 6, suggesting that the C6-derived RiPP may share similar functional and antimicrobial properties. Additionally, RiPPs and other secondary metabolites often undergo extensive post-translational modifications—including heterocyclization, oxidation, methylation, and glycosylation (Montalbán-López et al., 2021). Disruptions or variability in these modification processes can lead to end products that diverge from those predicted solely by genomic data (Arnison et al., 2013). A comparative analysis of RiPPs (ribosomally synthesized and post-translationally modified peptides) using Tanimoto similarity scores identified several peptides with closely related structural and biosynthetic characteristics. Five peptides Streptococcin A M49, Streptococcin A FF22, Salivaricin 9, Macedocin, and Ericin S exhibited a Tanimoto score of 1, indicating identical or nearly identical molecular fingerprints. These peptides belong predominantly to lanthipeptide class B and are produced by Streptococcus or Bacillus species, with biosynthetic modifications governed by LanM or LanBC enzyme systems. Additional peptides, including Lacticin 3147 A1, Subtilin, Entianin, Nisin Q, and Nisin F, also demonstrated high structural similarity, with Tanimoto scores ranging from 0.9375 to 0.9677. These peptides fall within lanthipeptide classes A and B and are biosynthesized by *Lactococcus lactis* or *Bacillus subtilis*, also utilizing LanM or LanBC modification systems. Overall, the findings underscore a strong degree of structural and biosynthetic conservation among lanthipeptides, particularly those modified via the LanM and LanBC enzymatic pathways, highlighting their evolutionary and functional relevance in antimicrobial activity.

**Table 6.** Structural Similarity of C6 Genome RiPP Precursor with Known Lanthipeptides.

| **RiPP** | **Tanimoto score** | **Class** | **Producer Organism** | **Modification System** |
| --- | --- | --- | --- | --- |
| <u>Streptococcin A M49</u> | 1 | lanthipeptideB | *Streptococcus pyogenes* | LanM |
| <u>Streptococcin A FF22</u> | 1 | lanthipeptideB | *Streptococcus pyogenes* | LanM |
| <u>Salivaricin 9</u> | 1 | lanthipeptideB | *Streptococcus salivarius* M18 |  |
| <u>Macedocin</u> | 1 | lanthipeptideB | *Streptococcus macedonicus* ACA-DC198 | LanM |
| <u>Ericin S</u> | 1 | lanthipeptideB | *Bacillus subtilis* | LanBC |
| <u>Lacticin 3147 A1</u> | 0.967742 | lanthipeptideB | *Lactococcus lactis* subsp. *lactis* | LanM |
| <u>Subtilin</u> | 0.966667 | lanthipeptideA | *Bacillus subtilis* | LanBC |
| <u>Entianin</u> | 0.966667 | lanthipeptideA | *Bacillus subtilis* subsp. *spizizenii* | LanBC |
| <u>Nisin Q</u> | 0.9375 | lanthipeptideA | *Lactococcus lactis* | LanBC |
| <u>Nisin F</u> | 0.9375 | lanthipeptideA | *Lactococcus lactis* | LanBC |

### 3.9. Molecular Interaction of antibiofilm Small molecules

Bacterial natural small biomolecules are soluble metabolites that are either released from lysed bacterial cells or secreted by live bacteria. These compounds, present in cell-free supernatants (CFS), exhibit notable biological activity and can exert specific physiological effects on the host. Several studies have reported that such metabolites possess antimicrobial and antibiofilm activities against *MRSA* and other pathogens (Khani et al., 2022; Narang et al., 2025; Yang et al., 2025). However, their precise mechanisms of action remain unclear. Therefore, in the present study, a literature mining approach was undertaken in July 2025 using search terms such as *“Lactobacillus plantarum and MRSA”* and *“cell-free supernatants, MRSA, and antibiofilm*.*”* A total of 15 potent antimicrobial and antibiofilm natural small biomolecules derived from the CFS of *L. plantarum* cultures were identified and selected for further investigation. To explore the anti-biofilm potential of *Lactobacillus plantarum*-derived 15 natural small biomolecules, molecular docking was conducted against Poly-β-1,6-N-acetyl-D-glucosamine synthase. It is a key enzyme encoded by the *icaA* gene of *Methicillin-resistant Staphylococcus aureus* (MRSA), critically involved in biofilm matrix synthesis (Nguyen et al., 2020; Iram et al., 2025). The binding affinities (in kcal/mol) of these biomolecules are depicted in Figure 1. Among the evaluated compounds, 2,4-Di-tert-butylphenol and Indole-3-lactic acid demonstrated the strongest binding affinities, with docking scores of −7.2 and −7.1 kcal/mol, respectively. These values suggest a high potential for effective binding with the target enzyme, indicating their significant inhibitory potential against biofilm formation. Similarly, Cyclo(L-propyl-L-valine) and 2,5-piperazinedione, 3,6-bis(2-methylpropyl) also exhibited strong interactions with the enzyme, with docking scores of −6.8 and −6.7 kcal/mol, respectively. Other noteworthy compounds include DL-4-Hydroxyphenyllactic acid and Pyrrolo[1,2-a] pyrazine-1,4-dione, both showing a moderate binding affinity of −6.4 kcal/mol. These molecules are known antimicrobial metabolites produced by *L. plantarum* and could potentially interfere with MRSA’s biofilm formation machinery through enzyme inhibition.

In contrast, simpler organic acids such as Acetic acid (−3.5 kcal/mol), Propionic acid (−4.1 kcal/mol), and Glyceric acid (−4.4 kcal/mol) displayed relatively weaker interactions, likely due to their smaller molecular size and limited functional groups for strong binding. However, their role should not be overlooked, as they may still contribute to the overall anti-MRSA activity through other synergistic or metabolic mechanisms. Lactic acid, Succinic acid, and L-Malic acid—typical fermentation end-products—showed moderate binding affinities (−4.5 to −5.6 kcal/mol), suggesting a possible contribution to biofilm inhibition, although their primary mechanism of action may be associated with environmental acidification rather than direct enzyme inhibition.

**Figure.**
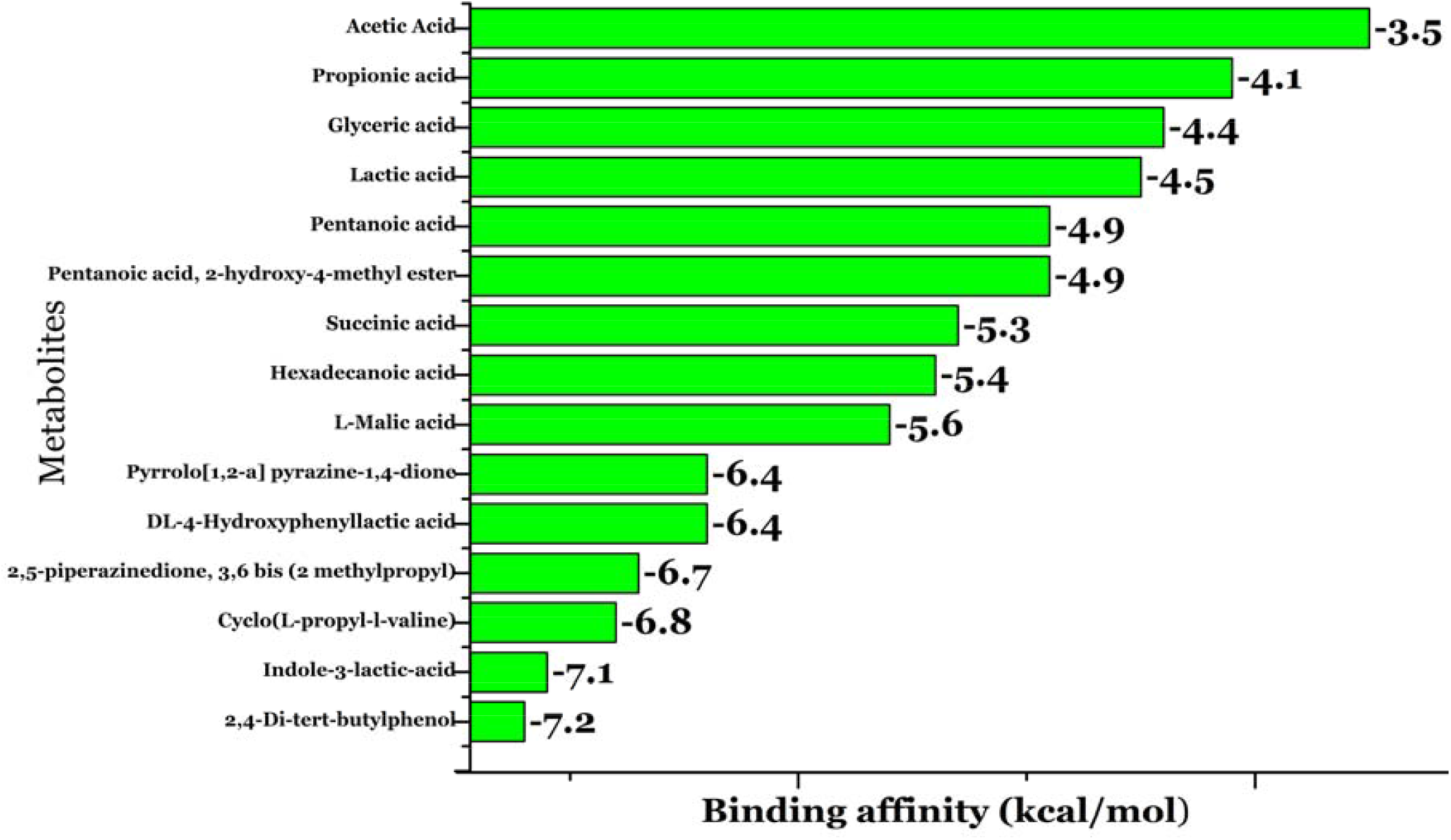
The bar graph illustrates the binding affinities (kcal/mol) of 15 metabolites from *L. plantarum* cell-free supernatant docked with the Poly-β-1,6-N-acetyl-D-glucosamine synthase. More negative values indicate stronger binding. Notably, 2,4-di-tert-butylphenol (–7.2 kcal/mol) and Indole-3-lactic acid (–7.1 kcal/mol) showed the strongest interactions, while Acetic acid (–3.5 kcal/mol) had the weakest.

**Figure.**
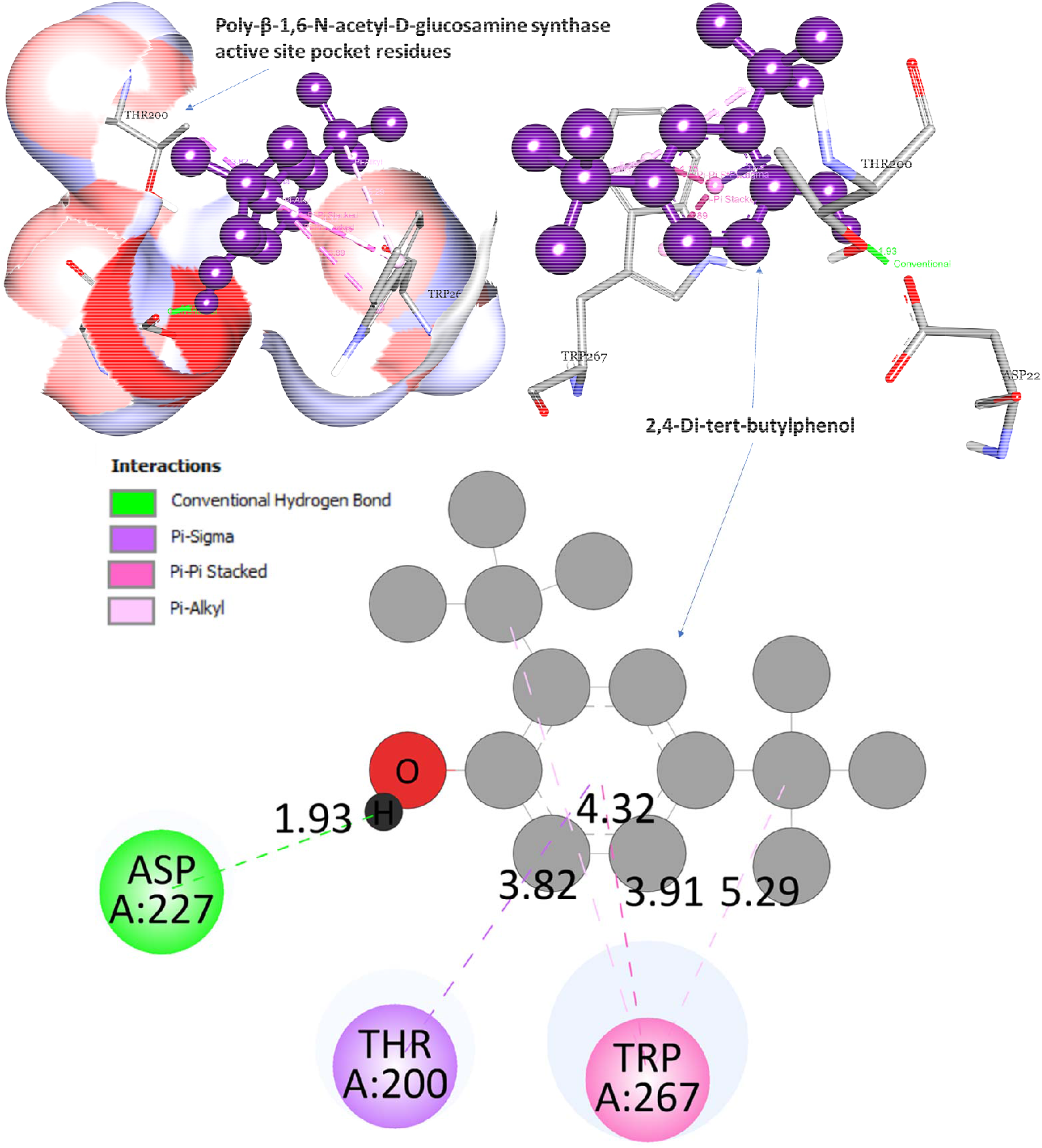
The interaction of 2,4-Di-tert-butylphenol with Poly-β-1,6-N-acetyl-D-glucosamine synthase invol strong non-covalent forces that stabilize its binding. A key hydrogen bond with ASP A:227 (1.93 Å) anch the ligand, while additional π–π, π–alkyl, and π–sigma interactions with TRP A:267 and THR A:200 enha hydrophobic stabilization. These interactions suggest the ligand fits well within the active site and m effectively inhibit the enzyme.

**Figure.**
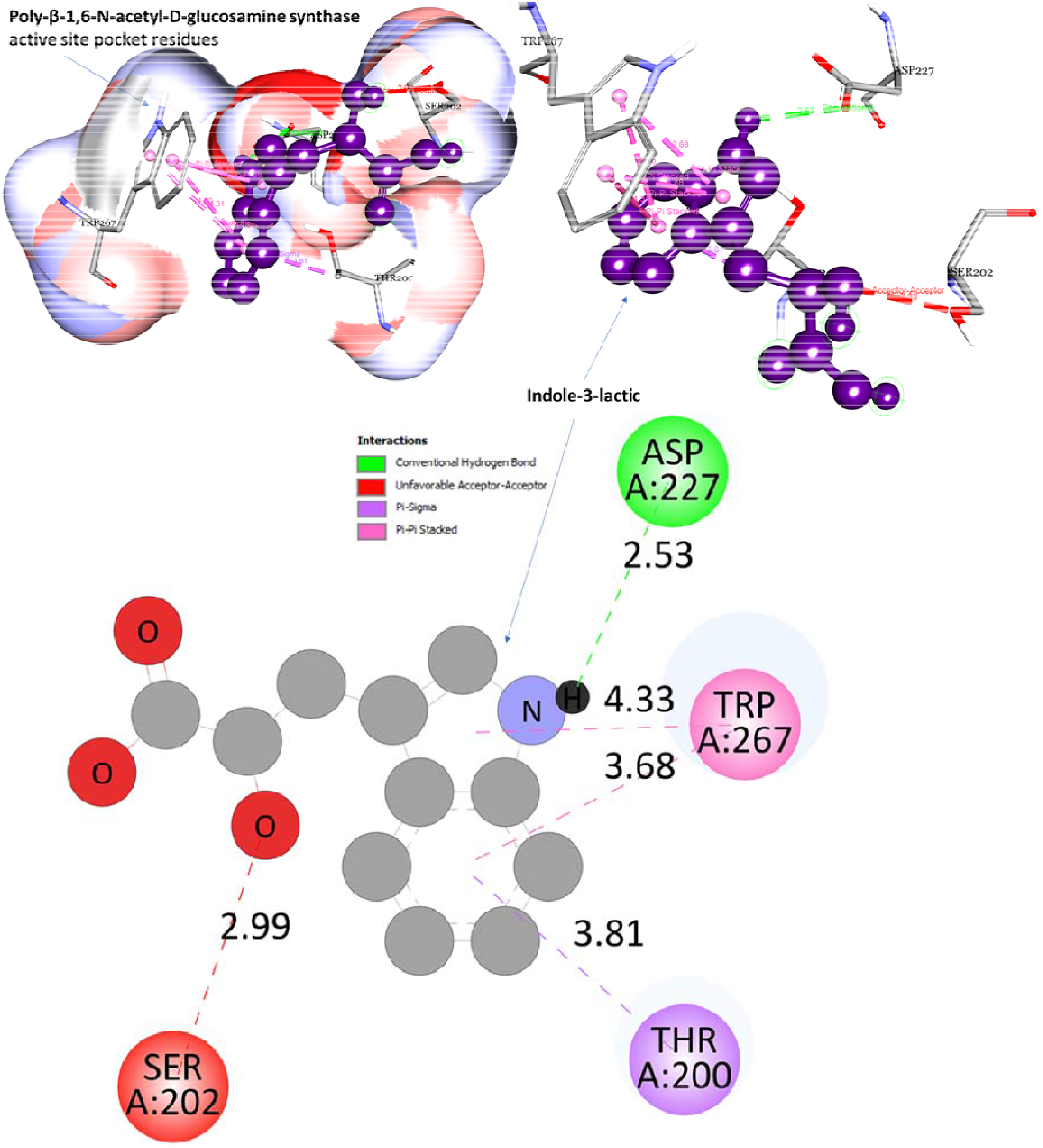
Indole-3-lactic acid binds moderately well to Poly-β-1,6-N-acetyl-D-glucosamine synthase through key interactions. A strong hydrogen bond with ASP A:227 (2.53 Å) stabilizes the ligand, while an unfavourable contact with SER A:202 (2.99 Å) suggests minor repulsion. Additional π–π, π–alkyl, and π–sigma interactions with TRP A:267 and THR A:200 further support binding through hydrophobic effects. These interactions indicate the ligand’s potential to influence enzyme activity.

Overall, the docking analysis suggests that specific secondary metabolites from *L. plantarum*, especially Indole-3-lactic acid and 2,4-Di-tert-butylphenol have high potential as inhibitors of Poly-β-1,6-N-acetyl-D-glucosamine synthase. These findings support the role of *L. plantarum* in disrupting MRSA biofilm formation and highlight its value as a source of postbiotic molecules for developing alternative anti-biofilm strategies.

## Conclusion

The comprehensive genome mining and *in silico* analysis of *Lactiplantibacillus plantarum C6* revealed the presence of multiple biosynthetic gene clusters (BGCs) associated with the production of diverse antimicrobial compounds, including RiPPs, lanthipeptides, terpenes, T3PKS, and cyclic-lactone inducers. Sequence similarity and structural analyses identified RiPP precursor peptides closely related to known bacteriocins, particularly from the lanthipeptide class, indicating a strong conservation of biosynthetic pathways. The presence of key regulatory genes within bacteriocin operons further supports the functional relevance of these clusters. Although experimental conditions may influence the detectability of some predicted peptides, the findings suggest that *L. plantarum* C6 possesses the genetic potential to synthesize a broad spectrum of bioactive secondary metabolites, underscoring its promise as a natural source of antimicrobial agents. Additionally, molecular docking studies demonstrated strong binding interactions between the predicted antimicrobial peptides and key target proteins from pathogenic organisms, supporting their potential functional role in antibiofilm activity. These docking results reinforce the bioactivity of genome-predicted peptides and highlight their possible application in inhibiting pathogen biofilm growth. The combined genome mining and docking data suggest that *L. plantarum* C6 harbors significant genetic and functional potential for the production of antimicrobial agents.

## DATA AVAILABILITY

The data that support the findings of this study are available from the corresponding author upon reasonable request.

## Supplementary material

The Supplementary Material for this article can be found online.

